# recount3: summaries and queries for large-scale RNA-seq expression and splicing

**DOI:** 10.1101/2021.05.21.445138

**Authors:** Christopher Wilks, Shijie C. Zheng, Feng Yong Chen, Rone Charles, Brad Solomon, Jonathan P. Ling, Eddie Luidy Imada, David Zhang, Lance Joseph, Jeffrey T. Leek, Andrew E. Jaffe, Abhinav Nellore, Leonardo Collado-Torres, Kasper D. Hansen, Ben Langmead

## Abstract

We present recount3, a resource consisting of over 750,000 publicly available human and mouse RNA sequencing (RNA-seq) samples uniformly processed by our new `Monorail` analysis pipeline. To facilitate access to the data, we provide the `recount3` and `snapcount` R/Bioconductor packages as well as complementary web resources. Using these tools, data can be downloaded as study-level summaries or queried for specific exon-exon junctions, genes, samples, or other features. `Monorail` can be used to process local and/or private data, allowing results to be directly compared to any study in recount3. Taken together, our tools help biologists maximize the utility of publicly available RNA-seq data, especially to improve their understanding of newly collected data. recount3 is available from http://rna.recount.bio.

## INTRODUCTION

RNA sequencing (RNA-seq) is a key tool in the study of disease and biology. The public Sequence Read Archive (SRA), which contains RNA-seq and other sequencing data types, doubles in size approximately every 18 months (Langmead and Nellore, 2018). We describe the recount3 project that makes archived RNA-seq datasets—both from the SRA and from compendia like The Genotype-Tissue Expression (GTEx) project—readily queryable, allowing users to access, combine and analyze datasets in new ways.

While other projects have summarized public RNA-seq datasets, most provide only gene- and transcript-level annotation-dependent summaries (Lachmann, Torre, et al., 2018; Ziemann et al., 2019; Tatlow and Piccolo, 2016). In addition to gene and transcript-level summaries, Toil produced annotation-guided splice junction quantifications (Vivian et al., 2017). The RNAseq-er gateway (Petryszak et al., 2017) continually analyzes RNA-seq datasets deposited in the European Nucleotide Archive and chiefly provides tabular gene and exon-level summaries, though genomewide annotation-agnostic coverage vectors are also available. The Expression Atlas (Papatheodorou et al., 2020) draws on datasets from GEO (Barrett et al., 2013) and Array Express (Athar et al., 2019), and the related Single Cell Expression Atlas (Papatheodorou et al., 2020) includes a further 180 single-cell RNA-seq studies from several species. But these contain only annotation-dependent gene-level summaries. Related work is detailed in Supplementary Note S1.

recount3 provides access to gene expression data in several resolutions and shapes, which enable studying expression using both gene reference annotation-dependent (gene, exon) and annotation-agnostic methods (exon-exon junctions, expressed regions) (Collado-Torres, Nellore, and Jaffe, 2017; Morillon and Gautheret, 2019). recount3 thus provides the basis for re-using public RNA-seq data to answer diverse biological questions.

recount3 includes a total of 316,443 human and 416,803 mouse run accessions (individual datasets) collected from the SRA, as well as large-scale human consortia including Genotype-Tissue Expression (GTEx version 8) and The Cancer Genome Atlas (TCGA). Second, recount3 includes several ways for users to query and use these annotation-dependent and annotation-agnostic expression summaries such as the `recount3` Bioconductor (Huber et al., 2015) package. The `snapcount` Biocon-ductor package, as well as the integrated Snaptron (Wilks et al., 2018) service, allow users to perform rapid queries across all summaries at once, e.g. across all the 316K human SRA samples. Lastly, our Snakemake-based (Köster and Rahmann, 2018) analysis pipeline `monorail` used to produce the summaries is designed to be easy for users to run on their own local RNA-seq reads across computing environments through the form of a single Docker/Singularity image.

Since its summaries were processed in a uniform and annotation-agnostic way, recount3 can fuel various analysis types. These could involve comparisons between human and mouse, cross-study comparisons, meta analyses, re-purposing of data to answer a new question, or broad explorations of the unannotated transcriptome. To illustrate, we survey splicing patterns in 700,000 run accessions and explore the fraction of exon-exon splice junctions that are present across several widely-used gene annotations. recount3 captures much of the cell type-specific splicing across various mouse cell types and cell type-specific junctions are depleted in gene annotations relative to junctions overall. We then demonstrate how our base-level coverage summaries reveal examples of non-coding and unannotated tissue- and cell-type specific transcription. Finally, we study the source of variation in gene expression across all the human and mouse samples, identifying key technical and biological sources of variation.

## RESULTS

### The recount3 resource

We developed a new distributed processing system for RNA-seq data called `Monorail` (Figure 1, Methods: `Monorail`, Supplementary Figures S1, S2, S3, S4, Supplementary Note S2). Using `Monorail`, we processed and summarized over 763K human and mouse sequencing runs, including GTEx v8, TCGA, and 732,246 runs from the Sequence Read Archive (Table 1), 416,803 of those from mouse. In compiling recount3, we processed 990 TB of compressed sequencing reads, used approximately 3.3 node-years of computation (29K node-hours) and produced over 150 TB of summarized data (Tables 1, 2, Supplementary Figures S5, S6, Supplementary Note S3).

**Table 1.** Size and computational cost of Monorail runs. Processing wall-clock times are estimated from run logs and are approximate. Wall times are roughly “node hours”, where a typical node used here has 48 cores and 192 GB of RAM. Node types vary somewhat across clusters used. Input and output sizes are calculated from compressed files. * Output size for GTEx v6 includes BAM files in addition to typical summaries † Number of additional junctions beyond those in GTEx v6. ‡ Total after counting only the distinct junctions.

| Collection | Input Size (TB) | Output Size (TB) | # Sequence Runs | # Studies | # Junctions (M) | # bigWigs (M) | Processing Wall Time (h) |
| --- | --- | --- | --- | --- | --- | --- | --- |
| SRA Human v3 | 474 | 72 | 316,443 | 8,677 | 228 | 1.2 | 1,728 |
| SRA Mouse v1 | 362 | 62 | 416,859 | 10,088 | 148 | 1.7 | 1,608 |
| TCGA | 75 | 7 | 11,348 | 1 | 31.5 | 0.045 | 170 |
| GTEx v6 | 35 | 6.7* | 9,911 | 1 | 22 | 0.040 | 168 |
| GTEx v7 & v8 | 46 | 4.9 | 9,303 | 1 | 10.6† | 0.037 | 123 |
| Total | 992 | 152.6 | 762,995 | 18,768 | 396‡ | 3.022 | 3,797 |

**Table 2.**
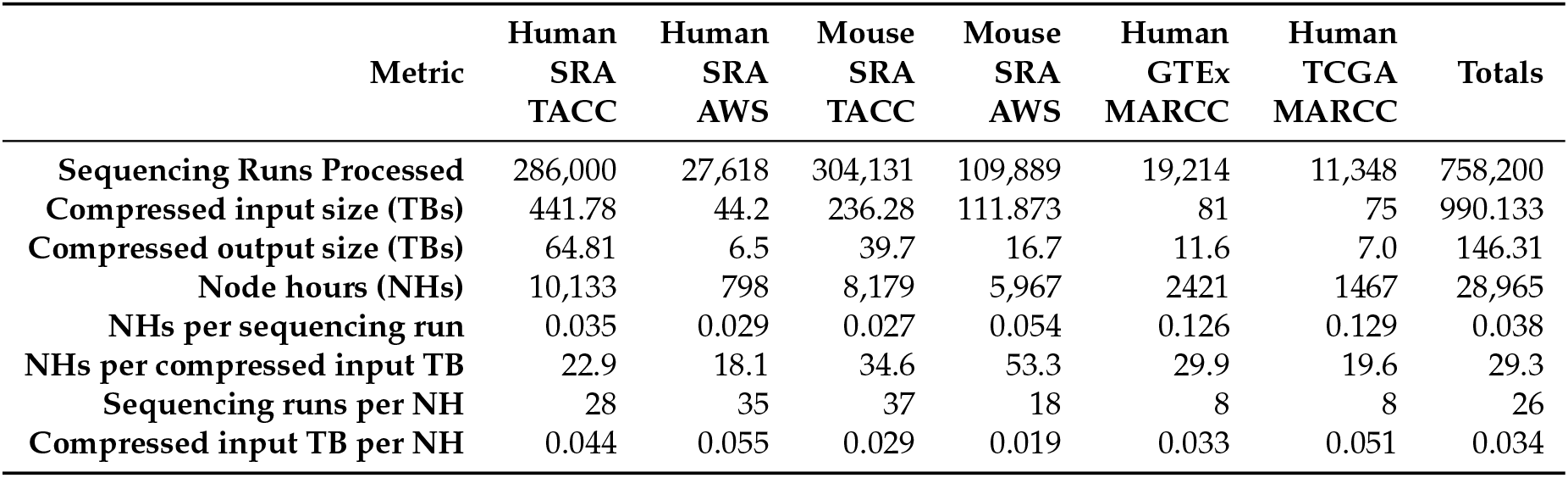
Monorail performance metrics run on TACC, AWS and MARCC. Statistics for GTEx and TCGA were extrapolated from a subset of each project (9277, 1567 samples respectively). GTEx output was increased by keeping whole BAM files for a subset of the samples. These numbers tally the number of run accessions processed, which can exceed the numbers in Table 1 due to some runs being processed multiple times, and due to runs that were later removed for QC or metadata reasons. Missing from this table are several thousand SRA human run accessions that were analyzed on MARCC but whose log files were discarded.

| Metric | Human SRA TACC | Human SRA AWS | Mouse SRA TACC | Mouse SRA AWS | Human GTEx MARCC | Human TCGA MARCC | Totals |
| --- | --- | --- | --- | --- | --- | --- | --- |
| Sequencing Runs Processed | 286,000 | 27,618 | 304,131 | 109,889 | 19,214 | 11,348 | 758,200 |
| Compressed input size (TBs) | 441.78 | 44.2 | 236.28 | 111.873 | 81 | 75 | 990.133 |
| Compressed output size (TBs) | 64.81 | 6.5 | 39.7 | 16.7 | 11.6 | 7.0 | 146.31 |
| Node hours (NHs) | 10,133 | 798 | 8,179 | 5,967 | 2421 | 1467 | 28,965 |
| NHs per sequencing run | 0.035 | 0.029 | 0.027 | 0.054 | 0.126 | 0.129 | 0.038 |
| NHs per compressed input TB | 22.9 | 18.1 | 34.6 | 53.3 | 29.9 | 19.6 | 29.3 |
| Sequencing runs per NH | 28 | 35 | 37 | 18 | 8 | 8 | 26 |
| Compressed input TB per NH | 0.044 | 0.055 | 0.029 | 0.019 | 0.033 | 0.051 | 0.034 |

**Figure 1.**
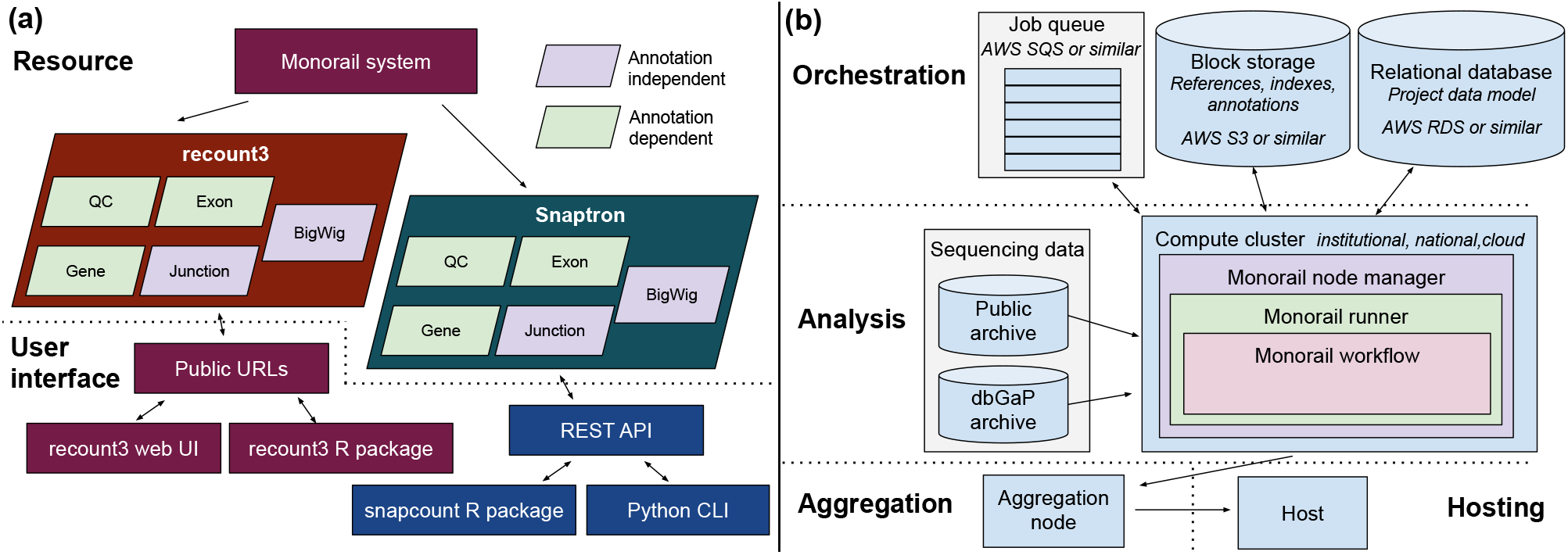
Overview of resource and grid design. **(a)** We use the `Monorail` system (detailed in panel b) to analyze and summarize data from archived sequencing datasets, yielding the recount3 and Snaptron resources. recount3 consists of five types of data summaries — Quality Control (QC), gene-level quantification, exon-level quantification, junction counts, and per-base coverages — packaged into tables with associated metadata and organized at the study level. The Snaptron resource consists of the same summaries, but organized at the level of a “collection,” where a single collection might include many studies, and indexed in a way that permits fast queries. Bottom: a few common ways that users can query and interact with the resources. QC, gene and exon-level summaries are compiled using gene annotations, but junction and bigWig coverage summaries are compiled without use of an annotation. **(b)** Illustration of `Monorail`’s grid design, split into Orchestration, Analysis and Aggregation layers. Supplementary Figures S1, S2, S3, S4 and Supplementary Note S2 contain details about `Monorail` and its analysis and aggregation components.

`Monorail` is not dependent on any specific compute backend and is capable of seamlessly utilizing heterogeneous compute resources. As an illustration of this, we highlight that recount3 was processed on 3 very different high performance systems: TACC, AWS and MARCC. TACC (Texas Advanced Computing Center) is part of the XSEDE super-computer ecosystem (Towns et al., 2014), AWS (Amazon Web Services) is a commercial cloud computing vendor and MARCC (Maryland Advanced Research Computing Center) is a traditional high-performance compute cluster. The `Monorail` system is available both as an open source suite of software, and as a self-contained public Docker image that produces identical results. This allows users to process private data and/or local read files using the same system used to produce recount3. In particular, `Monorail` can be used for processing dbGaP-protected data (Supplementary Note S4). Roughly half the runs in recount3 are from whole-transcript single-cell protocols such as Smart-seq (Goetz and Trimarchi, 2012) and Smart-seq2 (Picelli et al., 2013).

`Monorail` uses `STAR` (Dobin and Gingeras, 2016) and related tools to summarize expression at the gene and exons levels (annotation-dependent), to detect and report exon-exon splice junctions, and to summarize coverage along the genome as a bigWig file (Kent et al., 2010). The first step in this process consists of splicing-aware alignment of reads to the genome as well as quantification of exon-exon splice junction; this step is annotation-agnostic. Following this, as a second step, we quantify gene and exon expression using a specific annotation; for human samples we used Gencode v26, Gencode v29, FANTOM-CAT v6 and RefSeq v109 (Supplementary Table S1, Supplementary Note S5). The majority of the computational effort is spent on the annotation-agnostic first step. It is therefore possible to re-analyze the data using newer gene annotations or genomic regions of interest that use the same genomic coordinates, such as the FANTOM-CAT v6 case (Hon et al., 2017), with low computational effort. A detailed description of changes from recount2 (Collado-Torres, Nellore, Kammers, et al., 2017) is in Supplementary Note S6.

We developed a number of tools to facilitate user interaction with the processed data. Data can be accessed at the project/”dataset” level using the `recount3` R/Bioconductor package (Supplementary Note S7). We integrated processed recount3 data into the Snaptron (Wilks et al., 2018) system for indexing and querying data summaries. Further, we added a new R/Bioconductor interface to Snaptron called `snapcount` (Supplementary Note S8), which uses Snaptron to query recount3 summaries. Between the `recount3` and `snapcount` packages, it is easy to slice summarized data in various ways. For example, users can query specific genes across all samples or query specific projects across all genes. The results of these queries are delivered in annotated (across both samples/columns and genes/rows) `RangedSummarizedExperiment` objects (Huber et al., 2015). While we have focused on the R ecosystem, all our data are available in language independent data formats, including as text files and from the REST API from Snaptron.

Finally, we created a free, notebook-based computational resource where users can run R and Python based analyses on the same computer cluster at Johns Hopkins University where recount3 summaries are hosted. This makes use of the existing SciServer-Compute system (Taghizadeh-Popp et al., 2020) and allows users to run analyses on a free system where datasets are available locally, avoiding any extensive downloading. The contents of the resource and its user interfaces are illustrated in Figure 1a. In addition to SciServer-Compute, we have made recount3 available from AnVIL (Schatz et al., 2021), the genomic data science cloud platform from NHGRI.

In summary, we created an RNA-seq processing framework and resulting resource to facilitate reanalysis of hundreds of thousands of RNA-seq samples.

### Human and mouse splicing in SRA

Using recount3 splice-junction summaries, we surveyed unannotated splicing in the SRA, building on previous work (Nellore et al., 2016), but expanded to an unprecedented scale including both human and mouse. Further, we use an updated and expanded set of gene annotations, including Gencode (up to V33) and CHESS 2.2 (Pertea et al., 2018). We considered the subset of junctions that appear in at least 5% of SRA run accessions (15,773 out of 316,443 samples for human or 20,846 out of 416,803 for mouse, Methods: Analyses). We found that about 16% of human junctions and 12.5% of mouse junctions were not present in any tested annotation (Figure 2). Of the junctions in this subset, about 5% (human) and 3.5% (mouse) had both donor and acceptor sites present in the annotation, but not associated with each other, indicating an unannotated exon skipping or similar event. About 8.5% (human) and 7% (mouse) had either the donor or the acceptor present in the annotation, but not both. Remaining junctions (2.5% for human, 2% for mouse) had neither donor nor acceptor annotated. The 5% threshold is chosen to obtain junctions that might be considered “common”; we tested other thresholds in Supplementary Tables S2, S3.

**Figure 2.**
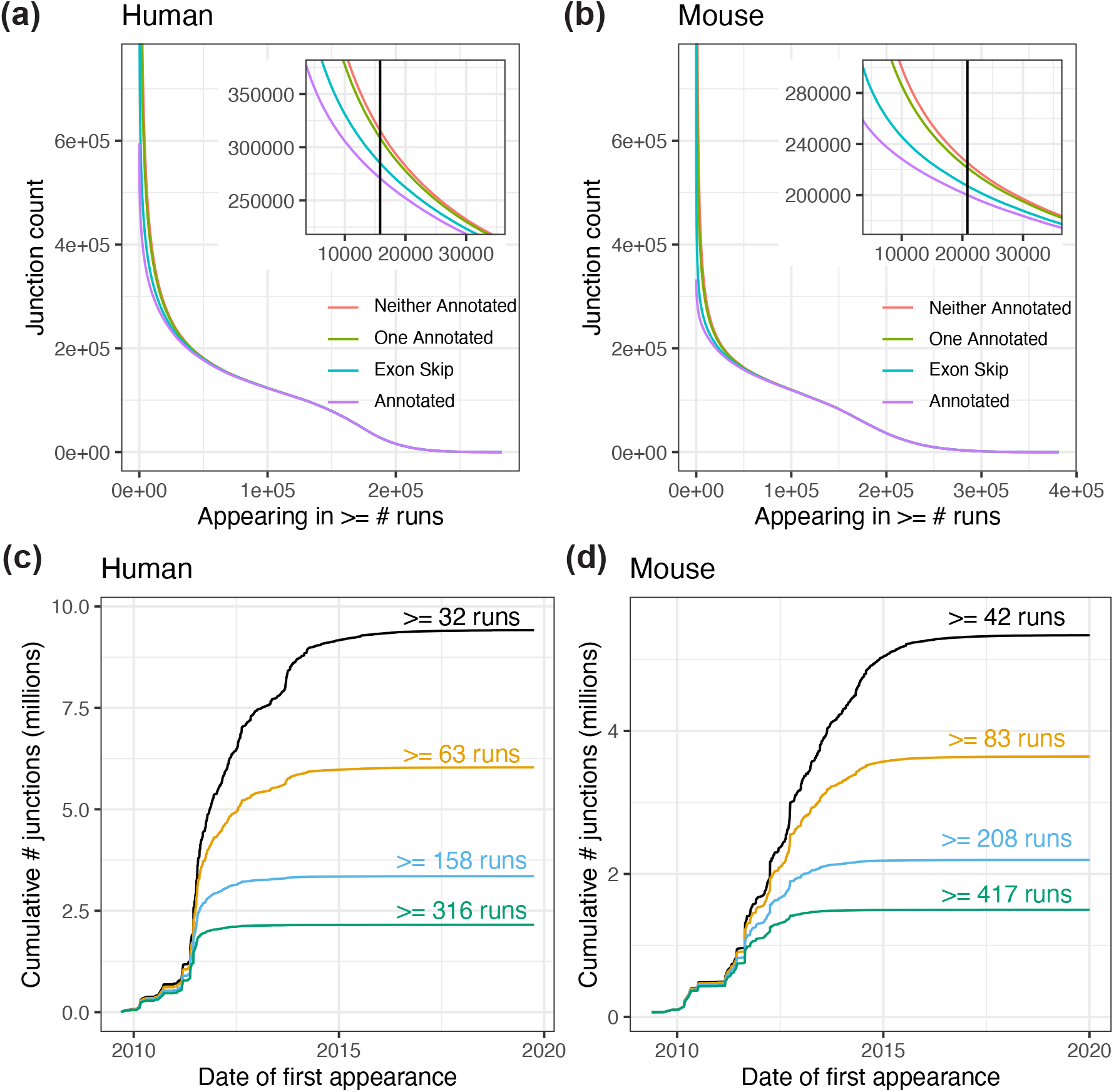
Annotation status and cumulative arrival of exon-exon splice junctions. **(a)** The number of exon-exon junctions present in at least *x* human runs accessions. Colors represent different degrees to which the junctions are annotated: completely annotated (purple) both donor and acceptor annotated but not in combination with each other (exon skip, blue), only donor or only acceptor annotated (one annotated, green), and neither annotated (red). Curves are cumulative across categories; thus, the number of unannotated junctions is given by the difference between the purple and red lines. **(b)** The same as (a) but for mouse run accessions. **(c)** The cumulative number of exon-exon junctions (in millions) across time for four different thresholds for the number of human run accessions where a junction is observed. **(d)** The same as (c) but for mouse run accessions.

We next asked whether cell type-specific splicing patterns tend to be annotated or unannotated. In the ASCOT study (Ling et al., 2020), we asked a similar question while focusing on cassette exons and on datasets where cell type was purified using fluorescence-activated cell sorting (FACS) or affinity purification. With recount3, we adapted this analysis to consider all splice junctions (not only cassette exons) and by additionally asking: what fraction of cell-type-specific splice junctions are present in any annotation? We considered the same purified datasets as the previous study, which included neuronal cell types, pancreas, muscle stem cells, CD4+ T-cells, B-cells, as well as ovary, testes, kidney, and stomach tissues among others. For each junction that occurred in at least one sample, we tested its cell type specificity using a Mann-Whitney U test comparing coverage within a cell type to coverage in all other cell types (403 samples in 34 studies). We binned the resulting −10 log p-values and calculated the percent of junctions in each bin that appeared in any tested gene annotation (Supplementary Table S1, Methods). We observed that more cell-type-specific junctions are less likely to appear in annotation (Figure 3). This suggests that the more specific a splicing pattern is to a particular cell type, the more likely it is to be ignored by analyses that quantify a gene annotation. These analyses highlight the utility of recount3 and producing RNA-seq alignments for splicing-based analyses of both annotated and unannotated sequences.

**Figure 3.**
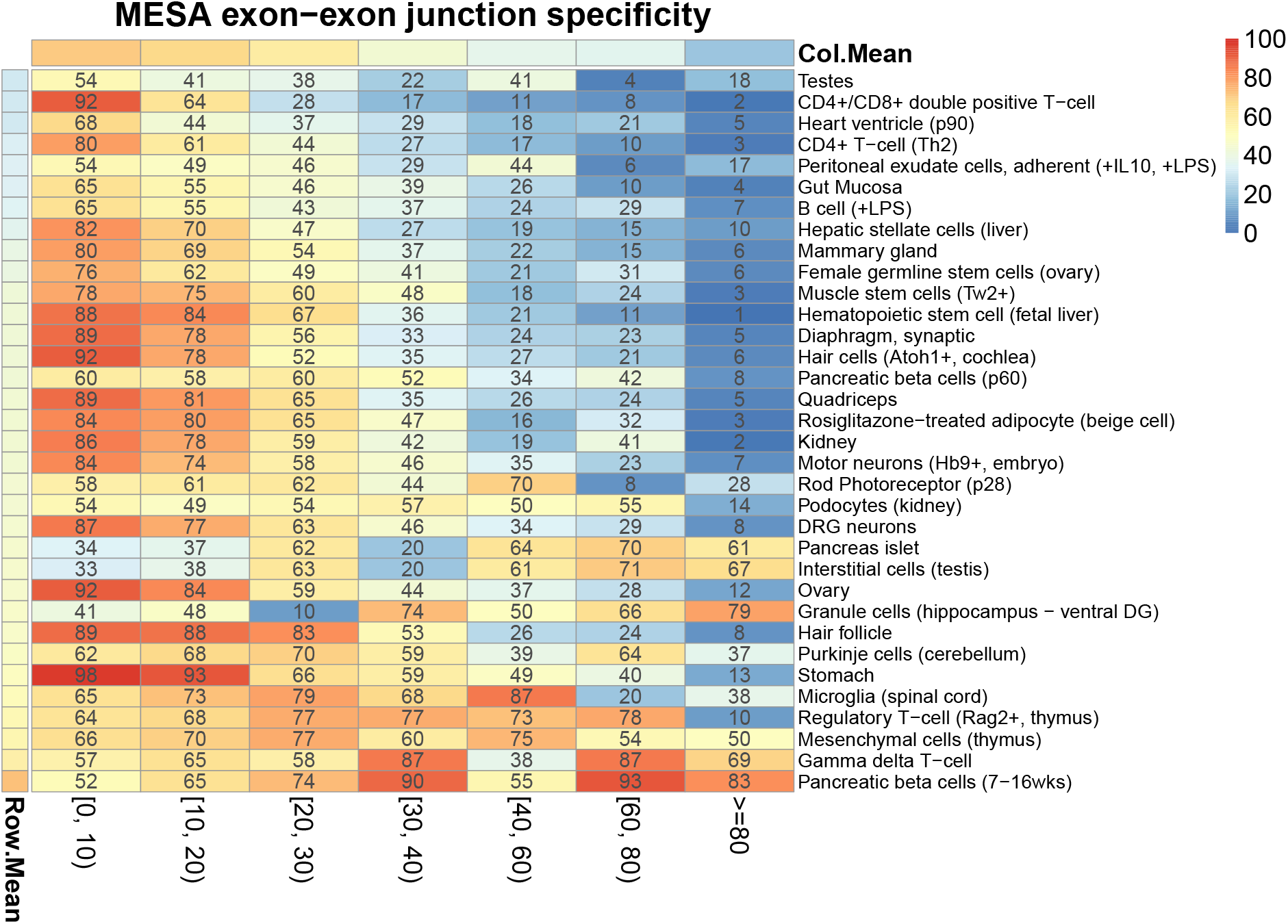
Cell-type enrichment of unannotated exon-exon junctions. Heatmap showing the cell-type enrichment results for novel exon-exon junctions similar to the one performed in the ASCOT study (Ling et al., 2020). The X-axis shows the bins of the −10 log p-values for cell type specificity using a Mann-Whitney U test comparing coverage within a cell type to coverage in all cell types. Each of the cell types evaluated are shown on the Y-axis with the percent of exon-exon junctions annotated denoted by a color gradient from dark blue (0%) to light yellow (50%) to dark red (100%). The row and column means are shown with the same color scale. The column mean, `Col.Mean`, shows that the average percent of annotated exon-exon junctions decreases as the cell type specificity increases on the X-axis.

### Non-coding and unannotated transcription

Since recount3’s bigWig files can be inputs to software for compiling gene and exon-level quantifications, we can quantify recount3 with respect to a new gene annotation that uses the same reference genome coordinates without re-aligning the reads. This is an important property of the recount3 resource as it enables quantifying expression summaries for diverse genomic regions of interest that annotation-dependent methods do not support, which can be done efficiently with `Megadepth` (Wilks et al., 2021). This feature facilitated generating the four quantifications included in recount3, ranging from smaller, more stringent annotations (RefSeq, O’Leary et al. (2016)), to more inclusive annotations (GENCODE, Frankish et al. (2019)), and to annotations focusing on 5’ boundaries and non-coding RNAs (FANTOM-CAT, Hon et al. (2017)).

The advantage of diverse annotations is illustrated by the FC-R2 study (Imada et al., 2020), which quantified recount2’s bigWigs using the FANTOM-CAT annotation, which includes a large number of non-coding RNAs (Hon et al., 2017). The study reported the tissue specificity of different classes of RNA: coding mRNA, divergent promoter lncRNA, intergenic promoter lncRNA, and enhancer lncRNA. Using recount3’s FANTOM-CAT quantifications, we updated that analysis to use the recount3 quantifications, including the additional runs present in GTEx v8 (FC-R2 used about half as many runs, Methods: Analyses). These results confirm those of the earlier study that ncRNAs tend to have more tissue-specific expression patterns than mRNA coding genes (Figure 4a).

**Figure 4.**
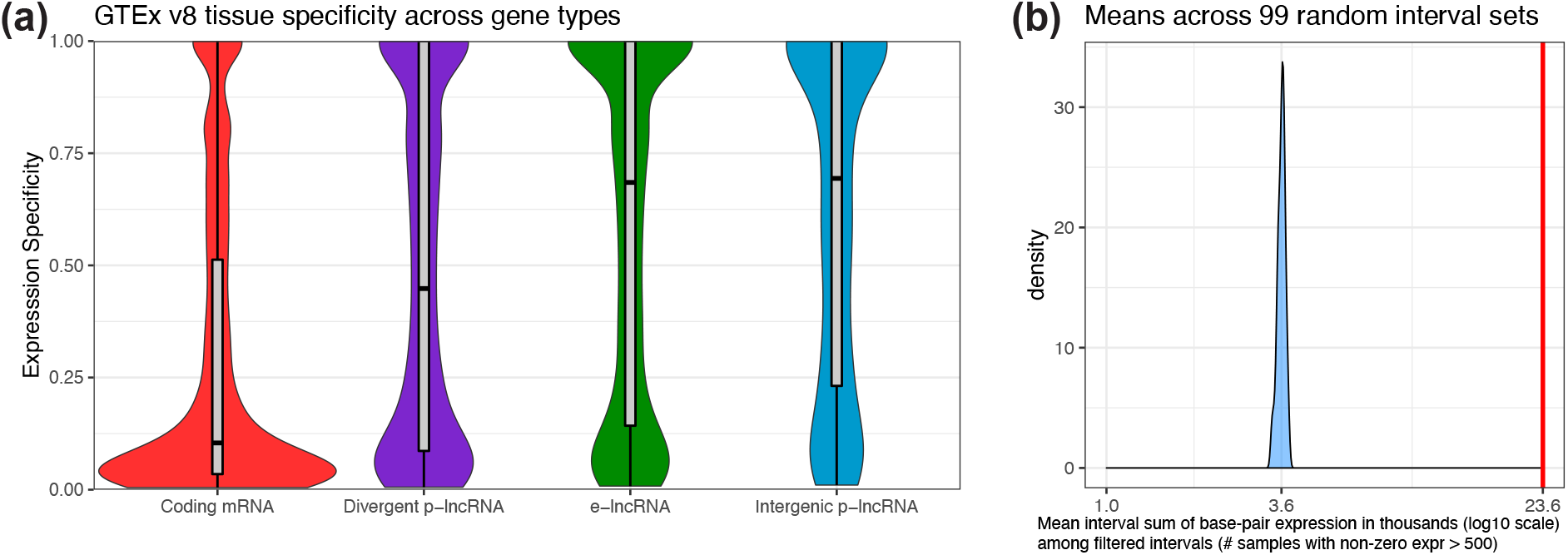
Non-coding and unannotated transcription. **(a) Expression profiles across GTEx v8 tissues.** Density and boxplots showing the tissue specificity of human coding and non-coding mRNAs from the FANTOM-CAT annotation (Imada et al., 2020) using data from GTEx v8. The specificity is based on the entropy computed from the median expression of each gene across the tissue types, as was done previously. **(b)** Evaluation of un-annotated genomic intervals powered by the base-pair coverage data in recount3. Previous work (Zhang et al., 2020) identified un-annotated genomic intervals that are expressed, conserved and overall not artifacts. Using the base-pair coverage data we can validate these genomic intervals compared to 99 sets of randomly selected intervals with the same length distributions matched by chromosome. This panel shows the density for the distribution, across the 99 random sets of intervals, of the mean sum of expression across base-pairs for intervals that are present in at least 500 samples. The red vertical line denotes the observed value from the intervals by Zhang et al. (2020).

To further show the utility of coverage-level summaries, we consider a recount2-based study by Zhang et al. (2020). With the premise that cell type-specific splicing patterns are less likely to be annotated, the authors used `derfinder` (Collado-Torres, Nellore, Frazee, et al., 2017) to analyze 41 GTEx v6 tissues and identify genomic intervals that were not present in any gene annotation but that were transcribed in a tissue-specific way. They found several such regions and used other sources of evidence (conservation, genetic constraint, protein coding ability) to argue that the discoveries could be potentially functional. Here we further use the bigWig files for the SRAv3 collection to show that the intronic ERs identified by Zhang et al have substantially more coverage in SRAv3 compared to length-matched, randomly chosen intronic intervals (Figure 4b, Methods: Analyses). The availability of coverage summaries therefore highlight the ease in re-quantifying RNA-seq alignments to updated and/or novel gene annotations and further permit the study of unannotated transcription.

### The landscape of transcription

To understand the sources of variation in gene expression, we conducted a principal component analysis (PCA) of the recount3 data, separately for human and mouse samples. One challenge in fully interpreting this analysis involves the incomplete sample metadata from SRA and GEO, so we manually curated a subset of studies including TCGA and GTEx (Methods: Analyses).

The largest source of variation for the expression of protein coding genes was whether the experiment type was bulk or single-cell (Supplementary Figure S7). We therefore built a simple predictor for experiment type (Methods: Analyses) and analyzed each type separately.

For bulk data, for both human and mouse, neither of the top principal components were correlated with library size (Figure 5a). Instead, the largest source of variation were correlated with tissue (cell) type (Figure 5b, Supplementary Figure S8). The clear clustering by tissue carries over to at least the first 4 principal components. In the human data, a large separate group consisted of blood samples (including lymphoblastoid cell lines, which are derived from B cells) (Figure 5c). Indeed, the first principal component of the human data is associated with the expression of hemoglobin genes (Supplementary Figure S9). Hemoglobin expression varies substantially even within blood samples because studies vary in the degree to which they deplete hemoglobin prior to assaying blood; GTEx is an example of a large study where hemoglobin was not depleted. These observations – including blood as an outlier group in the first two principal components for the human data – carry over to lncRNAs (Supplementary Figure S10.)

**Figure 5.**
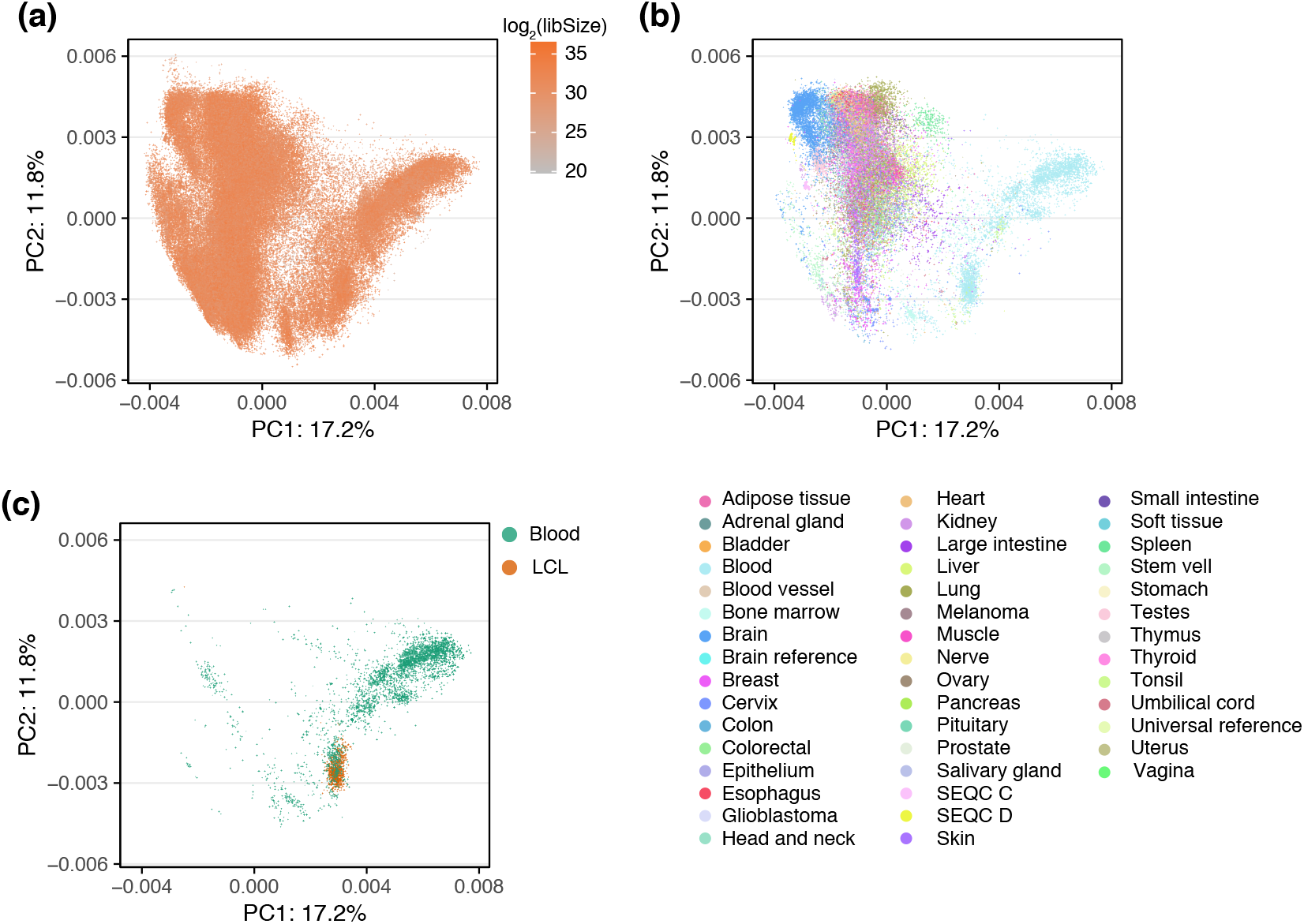
Gene expression variation. Principal component analysis of protein coding genes. **(a)** All human, bulk samples (165,880 samples) with color indicating the library size. **(b)** As (a) but only including samples with a labelled tissue (41,218 samples) colored by tissue. **(c)** As (a) but only including blood samples (4,731 samples) with color differentiating blood and lymphoblastoid cell lines (LCL) samples. Supplementary Figure S7 shows the variation across bulk and single-cell RNA-seq. The percentage of variance explained by the Principal Component (PC) is given in the axis labels.

For single-cell data, with far fewer studies available, and therefore a higher degree of confounding between tissue/cell type and study, the first principal component of the protein coding genes is library size, a technical covariate which is coupled to the degree of sparsity of the sample (Supplementary Figure S11, correlation between log_2_(library size) and the first principal component is 0.57). There is a singularity point with very low library size, where we observe very little between variation between samples. In contrast, as library size increases, the second principal component becomes cell type.

These large-scale analyses of gene expression levels can classify samples by experiment type and identify other important contributions of expression variation.

## DISCUSSION

recount3 is an easy-to-use resource for querying and obtaining annotation-dependent and annotation-agnostic expression summaries of public RNA-seq datasets. It includes roughly 800,000 run accessions divided evenly between human and mouse and includes all of GTEx v8 and TCGA. To create recount3, we created the `Monorail` system which runs on a variety of compute backends, including TACC/XSEDE, AWS and classic HPC systems. The availability of `Monorail`, as software and as a Docker image, makes it possible for users to process their own data for analysis alongside recount3 as described in the recount3 documentation website http://rna.recount.bio. This important `Monorail` feature enables comparing local and/or private data against the public data provided by recount3.

Importantly, the Monorail workflow does not use a gene annotation and is not biased against unannotated splicing patterns. While this requires that reads be aligned in a spliced fashion to the reference genome — a process that is more computationally expensive than quantifying a gene annotation — the resulting summaries enable crucial insights into the annotated transcriptome. Our results here showed that many well supported splicing events are unannotated and that cell-type-specific splicing events are particularly likely to be unannotatted, consistent with prior observations (Nellore et al., 2016; Ling et al., 2020). By further including base-pair resolution summaries, we also enable users to study transcription in any genomic region, annotated or not.

A strength of the Monorail workflow is that it can rapidly re-quantify datasets using new or updated gene annotations starting from the coverage (bigWig) files, bypassing the need to re-run the aligner or to store large BAM files. This will be important in the future as new annotations emerge and existing annotations undergo revisions and improvements.

We provide powerful interfaces via the `recount3` and `snapcount` R/Bioconductor packages, as well as through the Snaptron web service. Snaptron supports a wide variety of queries (eg. which samples supports a given splice-junction) and `snapcount` provides an R interface for these. Individual studies and metadata are accessible both through standard file formats (text, bigWig, Matrix Market) and through an R interface which delivers standard Bioconductor-supported objects for immediate analysis.

Though recount3 includes hundreds of thousands of run accessions, the utility of these data is often hampered by unreliable or missing metadata. This points to multiple directions for future work. First, it will be important to continue to build better models for predicting missing metadata and correcting mistakes in metadata (Ellis et al., 2018). Second, it will be important to enable users with more detailed knowledge of the datasets to create their own collections of related datasets, possibly with their own hand-curated metadata (Razmara et al., 2019), and allow the sharing of such hand-curated collections with the wider community. Third, since metadata can sometimes be an unreliable way to find relevant datasets, it will be important to design methods that search for related datasets based on their contents rather than their metadata, e.g. using genomic sketching (Ondov et al., 2016; Baker and Langmead, 2019).

A critical concern when combining public studies is technical confounding and batch effects (Leek et al., 2010). In light of this, it is perhaps surprising that our work shows the primary source of variation across all publicly available human and mouse RNA-seq data to be tissue, since we analyzed essentially uncorrected data. This agrees with previous work (Lee et al., 2020). However, we caution against over-interpreting this result. Here we studied total variation across all human and mouse tissues. In contrast, most biological questions of interest are more focused, concerning perturbations (such as disease) within a tissue or cell type. Expression variation associated with such perturbations is small compared to the total variation across all tissues. There is an ongoing need to adapt existing tools for removing unwanted variation to this setting of massive data with sometimes unreliable metadata, to make it possible to study such perturbations across data sources.

In summary, recount3 provides a way for all researchers, regardless of computing resources, to do large-scale analysis of gene expression.

## METHODS

### Monorail

#### Grid design

`Monorail`’s design follows the grid computing model. A large-scale analysis is centrally scheduled and orchestrated, with units of work handled by computers that might be spread across the world (Figure 1b). For our experiments, orchestration was handled by a collection of services running in the Amazon Web Services commercial cloud. Computing work was handled in three computing venues: (a) the Stampede2 cluster, located at the Texas Advanced Computing Center (TACC), accessed via the National Science Foundation’s XSEDE network, (b) instances on the AWS cloud’s Elastic Compute Cloud service, and (c) the Maryland Advanced Compute Center at Johns Hopkins University. The bulk of the work was performed at Stampede2, since it had both large capacity and no per-unit-time cost to us. But our grid design also enabled us to perform privacy-sensitive work on the local, dbGaP-approved MARCC cluster. Additionally, `Monorail` is designed to be fault tolerant, track provenance, centrally aggregate logs from all components of the architecture, and provide dashboards for monitoring and debugging. More details are provided in Supplementary Note S2.

#### Obtaining data

Sequence runs from both human and mouse were selected from the Sequence Read Archive (SRA) using filters designed to capture Illumina RNA-seq sequencing runs for bulk or whole-transcript single-cell sequencing experiments (Supplementary Table S4, Supplementary Note S9). We parsed metadata for these SRA runs and studies from the NCBI SRA metadata obtained using the Entrez API (Supplementary Note S9). For GTEx v8, we obtained the sequencing data from both SRA and the Google Cloud Platform, getting metadata from the from the Annotations section of the GTEx portal. For TCGA, we obtained the sequencing data using the GDC Download Client tool and inherited curated metadata from the earlier recount2 project (Supplementary Note S10).

#### Monorail: QC and Alignment

The `Monorail` analysis pipeline uses standard tools for analyzing RNA-seq data and compiling QC measures. In particular: (a) `STAR` (Dobin and Gingeras, 2016) to align RNA-seq reads in a spliced fashion to the reference genome, (b) `seqtk` (H Li, 2020) for analysis of base qualities and base composition, (c) `Megadepth` (Wilks et al., 2021) for analysis of fragment length and distribution of coverage across chromosomes, and (d) `featureCounts` (Liao et al., 2014) for a coarse QC analysis of gene expression levels. Details on the workflow used to generate expression and splicing summaries, as well as QC measures, can be found in Supplementary Note S11.

#### Monorail: Gene and exon quantification

`Monorail` uses `Megadepth` (Wilks et al., 2021) to analyze the spliced-alignment BAM file output by `STAR`. `Megadepth` produces a bigWig file representing depth of coverage at every genomic base (Supplementary Note S12). `Megadepth` also summarizes the amount of sequencing coverage within genomic intervals defined in a provided BED file. We build this BED file by taking all disjoint exonic intervals from one or more gene annotations. This allows us to quantify the intervals once using `Megadepth`, then sum the resulting quantities into exon- and gene-level quantities for all relevant annotations in the aggregation step. The specific gene annotations used are listed in Supplementary Note S5.

Given `Megadepth`’s coverage counts for the disjoint exonic intervals, `Monorail`’s aggregator uses a custom tool to sum these back into the full set of annotated genes and exons. The result is a tab-separated value (TSV) file containing coverage sums across all genes and exons in a set of gene annotations. This is the form the data takes in in recount3; being a TSV file, it is generically compatible with various programming languages and analysis environments. When the user obtains this data using the `recount3` R/Bioconductor package, the package translates the TSV file into a `RangedSummarizedExperiment` object (Huber et al., 2015) with all relevant row and column metadata. More information on the `recount3` R/Bioconductor package and the organization of the exon- and gene-level summaries is provided in Supplementary Note S7 and S13. See Supplementary Note S14 for information on the Snaptron data formatting and indexing steps.

#### Monorail: Summarizing splicing

When `STAR` performs spliced alignment, it outputs a high-confidence collection of splice-junction calls in a file named (`SJ.out.tab`). This file describes each junction called by `STAR`, including its beginning and end, strand, splice motifs, and the number of spliced alignments supporting the junction call. Starting from these files — one per run accession — the `Monorail` aggregator combines them into both study- and collection-level matrices. These matrices are naturally sparse, having mostly 0 entries, since many splice junctions appear only in a small number of run accessions. Study-level junction-count matrices are then converted into Matrix Market and included in the recount3 resource. Collection-level matrices are indexed using SQLite and/or tabix in preparation for Snaptron queries.

#### Running Monorail

We used Stampede2, AWS EC2, and an institutional cluster (MARCC) to process approximately 760,000 human and mouse sequencing runs comprising 990 TB of compressed data over six months starting October 2019 (Table 2). We used about 29,000 node hours in total, or 0.038 node-hours per sequencing run. We estimate this would cost about $0.037 per accession using equivalent cloud resources, improving substantially on the $0.91 per accession achieved by our previous `Rail-RNA` system (Nellore et al., 2017). Details of the cost calculation are given in Supplementary Note S3.

Roughly speaking, this cost is higher but within a factor of about four times the per-accession costs of other large-scale analysis pipelines (Ziemann et al., 2019; Lachmann, Torre, et al., 2018). The difference is because `Monorail` produces a wider palette of outputs — e.g. a summary of both annotated and unannotated splicing, bigWig files — allowing more downstream analyses and relying less on a gene annotation. Also, other systems generally require a re-run of the workflow to adapt to a change in gene annotation; `Monorail` can quantify a new gene annotation directly from the per-base coverage files, bypassing realignment. Details on `Monorail` cost and throughput are provided in Supplementary Note S3.

#### Analyses

The GitHub repositories with the code for these analyses are listed at http://rna.recount.bio/docs/related-analyses.html.

#### Human and mouse splicing in SRA

We adapted the analysis from the ASCOT study (Ling et al., 2020), which gathered various studies of purified cell types in mouse, to consider all splice junctions individually rather than as cassette exons. Given the junction-level summaries produced by the Monorail workflow, we measured the fraction of cell-type-specific splice junctions present in any of recount3’s gene annotations for mouse. For each junction that occurred in at least one sample, we tested its cell type specificity using a Mann-Whitney U test comparing coverage within a cell type to coverage in all other cell types (403 samples in 34 studies). We binned the resulting −10 log p-values and calculated the percent of junctions in each bin that appeared in any tested gene annotation (Supplementary Table S1).

#### Non-coding and unannotated transcription

Using recount3’s FANTOM-CAT quantifications, we updated the previous analysis (Imada et al., 2020) to use the recount3 quantifications, including the additional runs present in GTEx v8. Briefly, this involving computing the counts per million (CPM) for the GTEx v8 data, then computing the entropy across the different GTEx tissues, and using the FANTOM-CAT annotation (Imada et al., 2020) we made Figure 4a.

We obtained the intronic ERs identified previously (Zhang et al., 2020), separated them by chromosome, to then generate randomly chosen intervals with the same length distribution from the same chromosomes as the intronic ERs. We repeated this process 99 times in order to generate 99 sets of random intervals that are length and chromosome-matched to the intronic ERs. We then quantified the sum of the base-pair coverage across each of those random intervals on the human bigWig files from SRA studies using `Megadepth` version 1.0.3 (Wilks et al., 2021) with the `--annotation` and `--op sum` arguments. We then filtered the random intervals to keep those with non-zero expression in at least 500 samples. Then we computed the mean sum of base-pair expression among the random intervals for each of the 99 random sets, and finally we show the distribution of these 99 means in Figure 4b.

#### Differentiating between bulk and single-cell

We built a simple predictor based on the sparsity pattern of a set of labelled data. We estimate empirical distributions of the percentage of genes with 0 expression, separately for each of the two labelled groups, yielding two observed training distributions *f*_*b*_ and *f*_*sc*_. Samples were predicted to be either bulk or single-cell based on the ratio *r* = *f*_*b*_/(*f*_*b*_ + *f*_*sc*_). We also considered using library size as predictor, but decided to not include it.

For the mouse data we used simple text analysis (Supplementary Note S9) of the SRA meta data records to predict whether a sample was single-cell or bulk; this was used as labelled data. We made predictions on all samples (which could differ from the predictions based on text analysis), using the cutoffs *r*(sample) *>* 0.825 to predict bulk, *r*(sample) *<* 0.1 to predict single-cell and the remaining samples were unclassified. We used the gencode v25 annotation.

For the human data we used a larger set of labelled samples, both based on the text analysis (as for mouse) and also manual curation. We had *∼*50k manually curated samples (of which *∼*48k was bulk) and *∼*92k samples based on text analysis. Predictions were made using the cutoffs *r*(sample) *>* 0.5 to predict bulk and *r*(sample) *<* 0.5 to predict single-cell. We used the gencode v29 annotation.

#### Manual curation

We performed PCA on the protein coding genes in mouse to investigate gene expression variation. Using both the PCA plot as well as study size, we chose studies for manual curation, based on selecting 2-4 studies out of the 5 largest studies. We subsequently added samples from studies of testes.

For the human data we manually curated tissue types and experiment types for 30,473 SRA samples of 373 SRA studies. We used the meta-data provided by SRA selector or corresponding GEO samples. The experiment types are broadly classified into 4 categories: bulk RNA-seq, single cell RNA-seq, small/Micro RNA-seq, and others. Others consist of a range of experiment types, such as locus-targeted sequencing and ribosome sequencing, which are usually quite different from traditional bulk RNA-seq (Supplementary Figure S7); users should check the SRA database if interested in these other experiment types. We supplemented these SRA studies with GTEx and TCGA.

### Data Presentation

#### recount3

The `recount3` R/Bioconductor package allows users to download gene, exon, and exon-exon junction counts data provided by the recount3 resource. *recount3* is designed to be user friendly and enable users to utilize the full set of analytical and visualization software available in Bioconductor for RNA-seq data. `recount3` achieves this by enabling downloads per study and by presenting the data through `RangedSummarizedExperiment` R/Bioconductor objects (Huber et al., 2015). `recount3` provides multiple options for converting the base-pair coverage counts (Collado-Torres, Nellore, and Jaffe, 2017) into read counts, RPKM values, among other options. Furthermore, `recount3` provides the URLs for accessing all of the recount3 resource text files such as the sample bigWig coverage files (Kent et al., 2010), enabling non-R users to build their own utilities for accessing the data. See Supplementary Note S7 for more details.

#### Snaptron

While `recount3` offers the user a way of accessing gene, exon, and junction coverage, it is limited to providing that only at the study level. Snaptron (Wilks et al., 2018) and its newly added R interface, `snapcount`, provide the ability to query precise regions of the genome for the coverage generated in `Monorail`. Queries can be made across all samples or for a specific subset. Queries can request summaries at the gene, exon, or junction level. Queries can be further filtered by aggregate sample occurrence and read coverage. Additionally, these tools enable “higher-level” analyses to be carried out across region queries to support operations such as percent spliced in (PSI) and tissue specificity (in the case of GTEx). `snapcount` specifically creates filtered `RangedSummarizedExperiment` objects (Huber et al., 2015) dynamically based on the user’s query in contrast to the fixed nature of recount3’s study-level data objects. See Supplementary Note S8 for more details.

## SUPPLEMENTARY MATERIALS

### SUPPLEMENTARY TABLES

**Supplementary Table S1.**
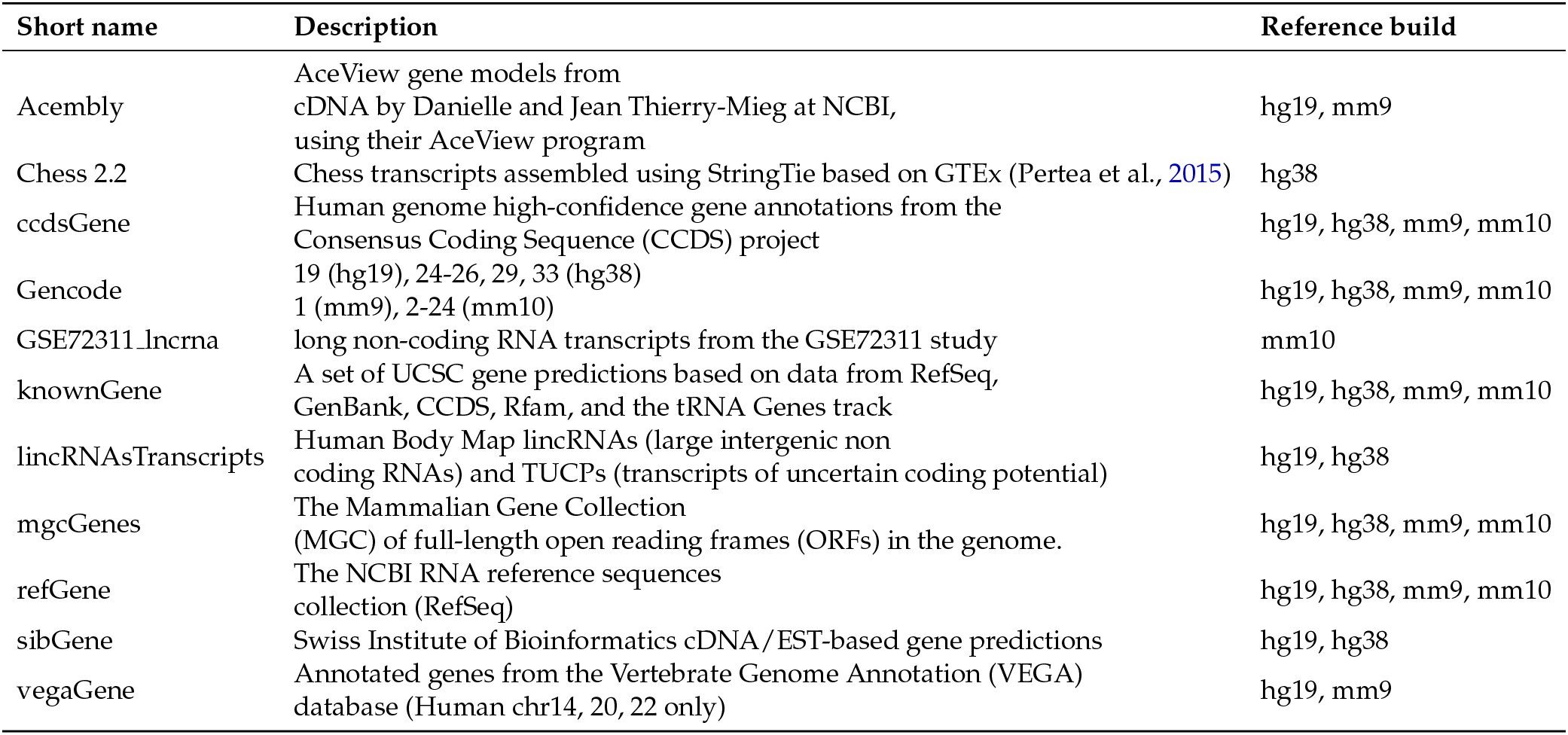
Junction annotation sources. Descriptions are from the UCSC Table Browser track detail page or the Gencode website. See also Supplementary Note S5.

**Supplementary Table S2.**
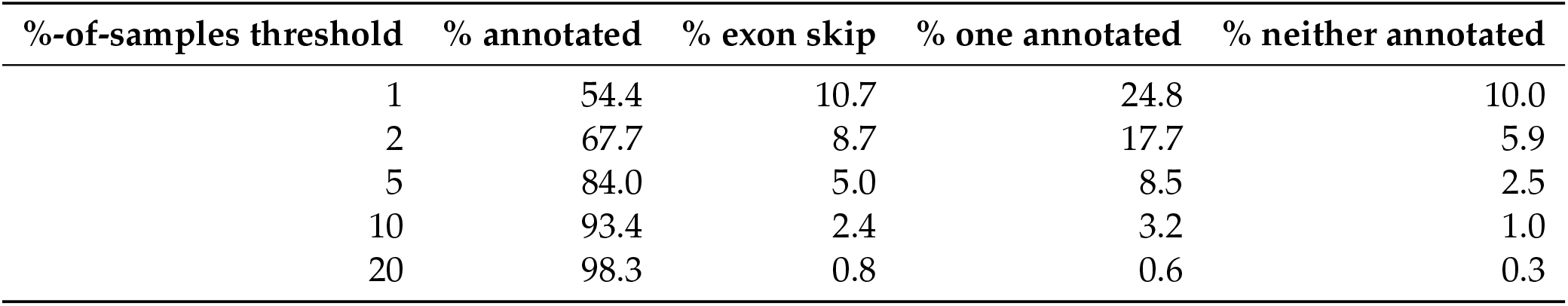
SRA Human v3 Annotated Junction Percentages. First column specifies the filtering criterion: only junctions appearing in at least this percent of run accessions are included. Other columns specify the percentage of these junctions that appear entirely in some annotation (“% annotated”), where both the donor and acceptor appear in some annotation, but never together in the same junction (“% exon skip”), where either the donor or the acceptor do not appear in annotation (“% one annotated”), and where neither donor nor acceptor appear in annotation (“% neither annotated”). We used an inclusive set of gene annotations including all the annotated junctions in several Gencode versions up to V33, as well as CHESS 2.2. The full set of annotations is listed in Supplementary Table S1.

| %-of-samples threshold | % annotated | % exon skip | % one annotated | % neither annotated |
| --- | --- | --- | --- | --- |
| 1 | 54.4 | 10.7 | 24.8 | 10.0 |
| 2 | 67.7 | 8.7 | 17.7 | 5.9 |
| 5 | 84.0 | 5.0 | 8.5 | 2.5 |
| 10 | 93.4 | 2.4 | 3.2 | 1.0 |
| 20 | 98.3 | 0.8 | 0.6 | 0.3 |

**Supplementary Table S3.** SRA Mouse v1 Annotated Junction Percentages. Same as previous table, but considering junctions from SRA Mouse v1. We used an inclusive set of gene annotations including all the annotated junctions in Gencode versions up to V24. The full set of annotations is listed in Supplementary Table S1.

| %-of-samples threshold | % annotated | % exon skip | % one annotated | % neither annotated |
| --- | --- | --- | --- | --- |
| 1 | 55.7 | 8.7 | 23.9 | 11.6 |
| 2 | 70.6 | 6.8 | 16.5 | 6.2 |
| 5 | 87.5 | 3.5 | 7.0 | 2.0 |
| 10 | 95.4 | 1.5 | 2.4 | 0.7 |
| 20 | 98.8 | 0.5 | 0.4 | 0.2 |

**Supplementary Table S4.** Bulk versus single-cell run accessions processed. Metadata fields were used to subdivide the run accessions into “Smart-seq” and “Bulk” categories, according to criteria described in Supplementary Note S9.

| | | Runs | Studies | Bases<br>$\times 10^{12}$ | Terabytes<br>(compressed) |
| --- | --- | --- | --- | --- | --- |
| **Human** | Bulk | 218,978 | 8,357 | 868 | 451 |
|  | Smart-seq | 97,465 | 320 | 60 | 30 |
|  | Total | 316,443 | 8,677 | 928 | 481 |
| **Mouse** | Bulk | 203,144 | 9,407 | 657 | 329 |
|  | Smart-seq | 213,715 | 681 | 75 | 34 |
|  | Total | 416,859 | 10,088 | 732 | 363 |

### SUPPLEMENTARY FIGURES

**Supplementary Figure S1.**
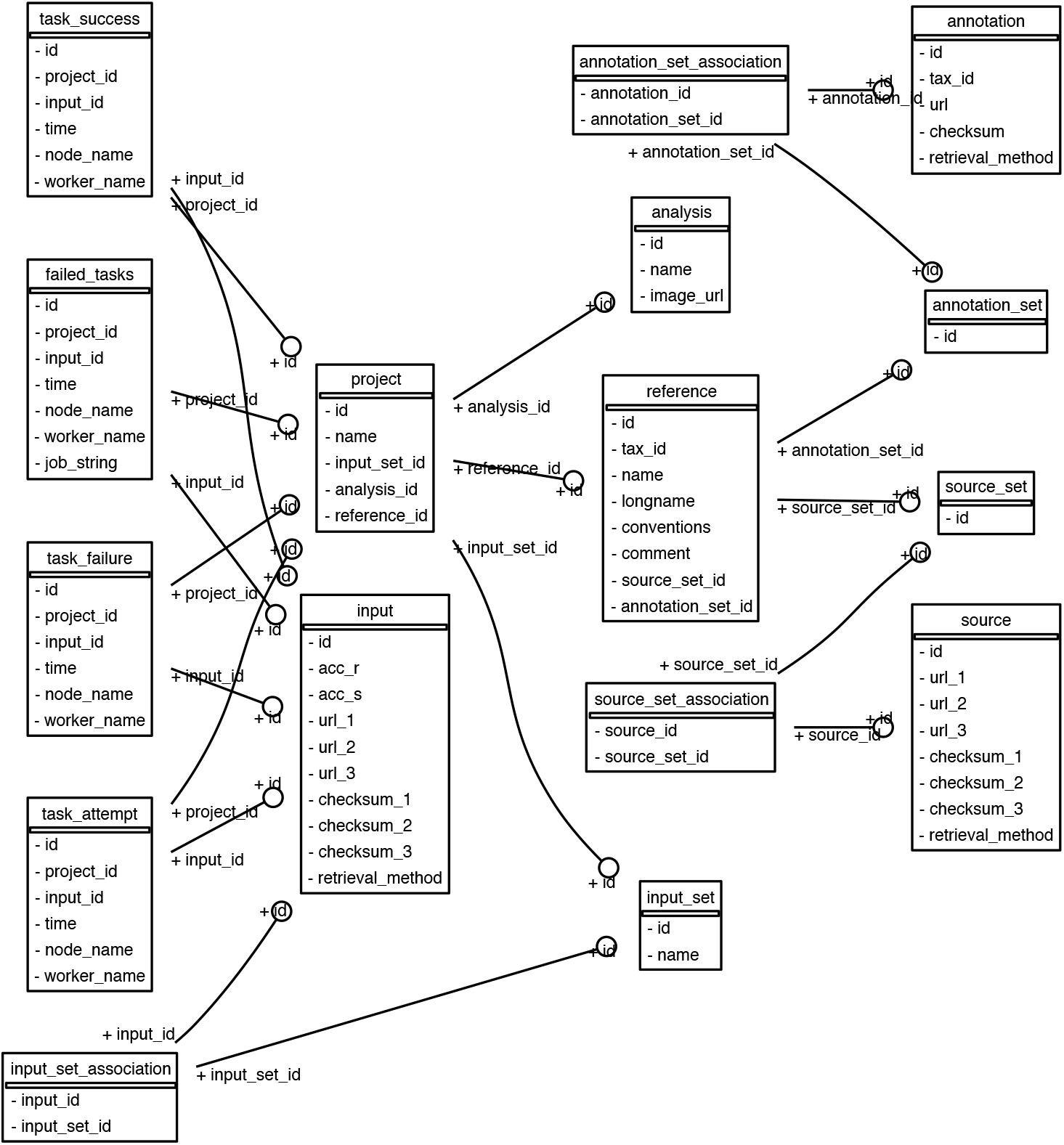
The `Monorail` relational database model. Rectangles denote tables and arcs denote the key relationships between tables. This image was created using the `sqlalchemy schemadisplay` package. See also Figure 1 and Supplementary Note S2.

**Supplementary Figure S2.**
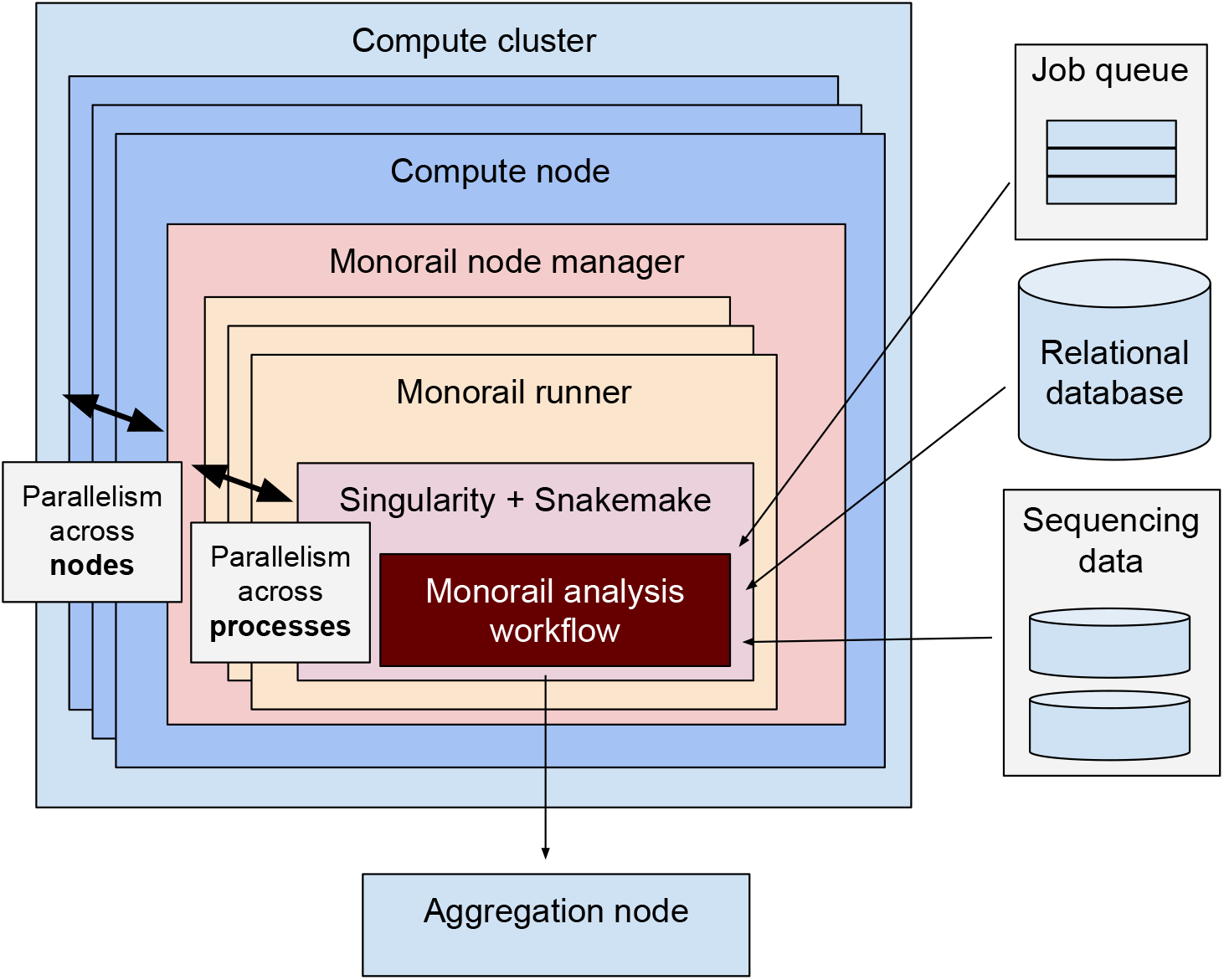
Monorail workflow parallelism. Illustration of how `Monorail` ensures parallelism both across nodes of the cluster (node manager launching many simultaneous runners) and across cores on a single cluster node (runner launching many simultaneous processes). Supplementary Figure S3 shows a single `Monorail` analysis workflow. See also Figure 1 and Supplementary Note S2.

**Supplementary Figure S3.**
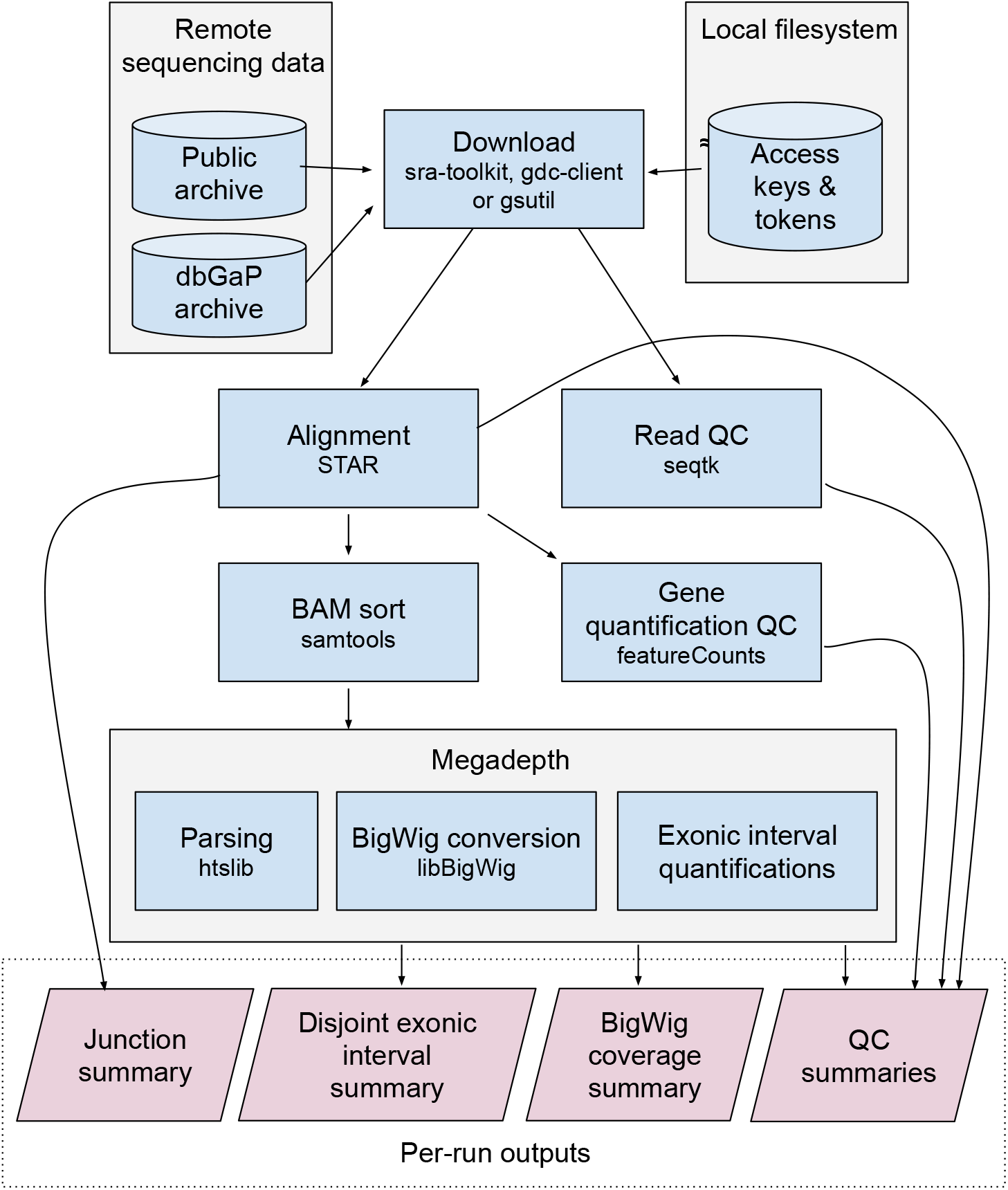
Monorail analysis workflow. Illustration of key steps and key inputs and outputs of the `Monorail` analysis workflow. Not shown: reference, gene annotation and index files are also stored on the local filesystem and loaded by tools like `STAR` and `featureCounts` as needed. The workflow is driven by Snakemake and runs within a Singularity container on a cluster node. Supplementary Figure S2 illustrates how many such workflows run in parallel on a cluster. See also Figure 1 and Supplementary Note S2.

**Supplementary Figure S4.**
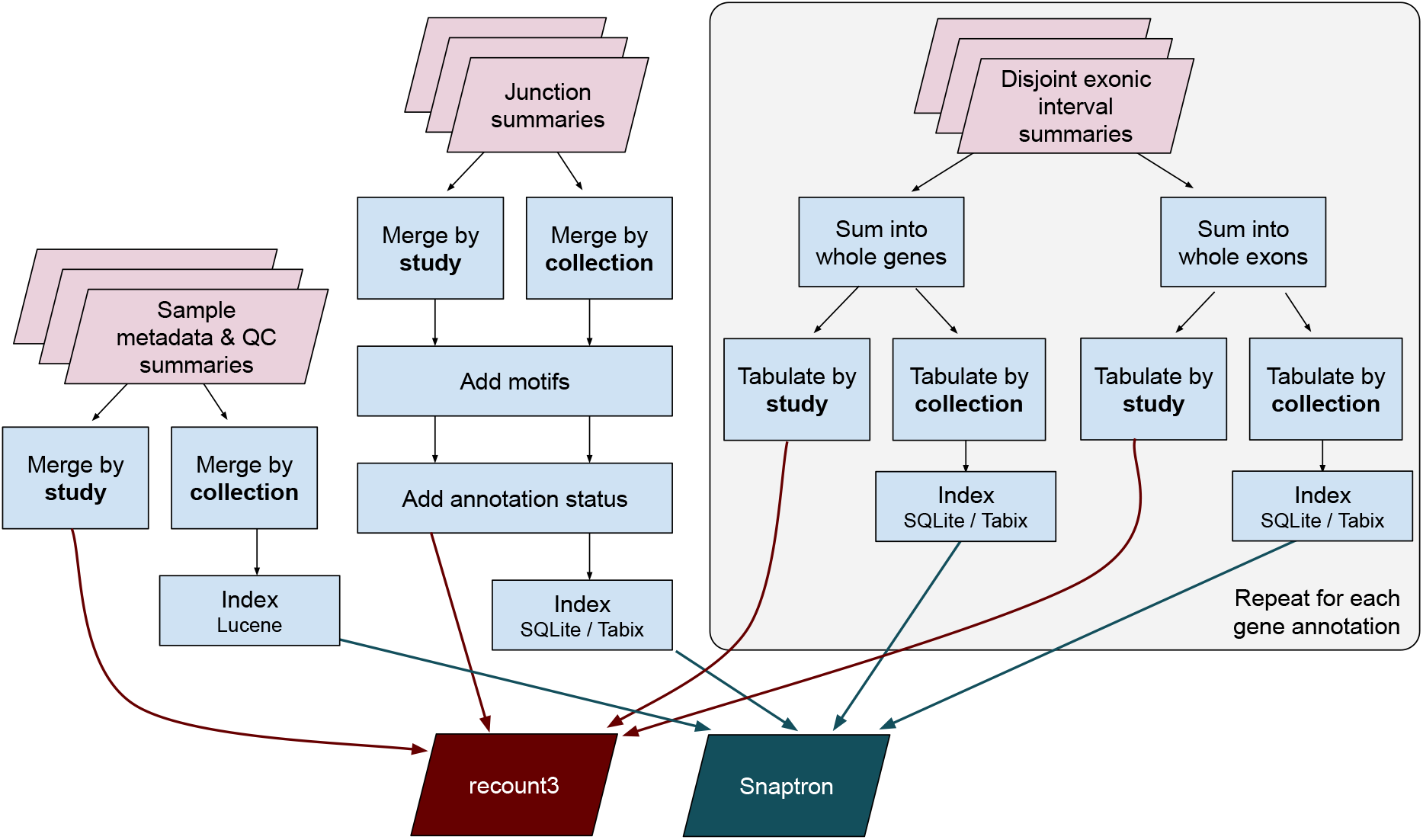
Monorail Aggregation Workflow. Illustration of `Monorail`’s aggregation workflow. The workflow runs on a single computer, taking all run-level analysis outputs as its input. Its outputs are the study- and collection-level objects and indexes hosted by recount3 and Snaptron. The disjoint exonic interval summaries furnish raw counts that can be summed into both exon- and gene-level counts. The “add motifs” step for junctions adds the specific donor and acceptor dinucleotide patterns present in the reference in genome for that junction. For the “add annotation status” step for junctions, an inclusive set of gene annotations is used, as detailed in Supplementary Note S5. Not pictured: metadata is also aggregated and hosted by recount3 and Snaptron. Also not pictured: bigWig files are organized into per-study subdirectories on the host filesystem. See also Figure 1 and Supplementary Note S2.

**Supplementary Figure S5.**
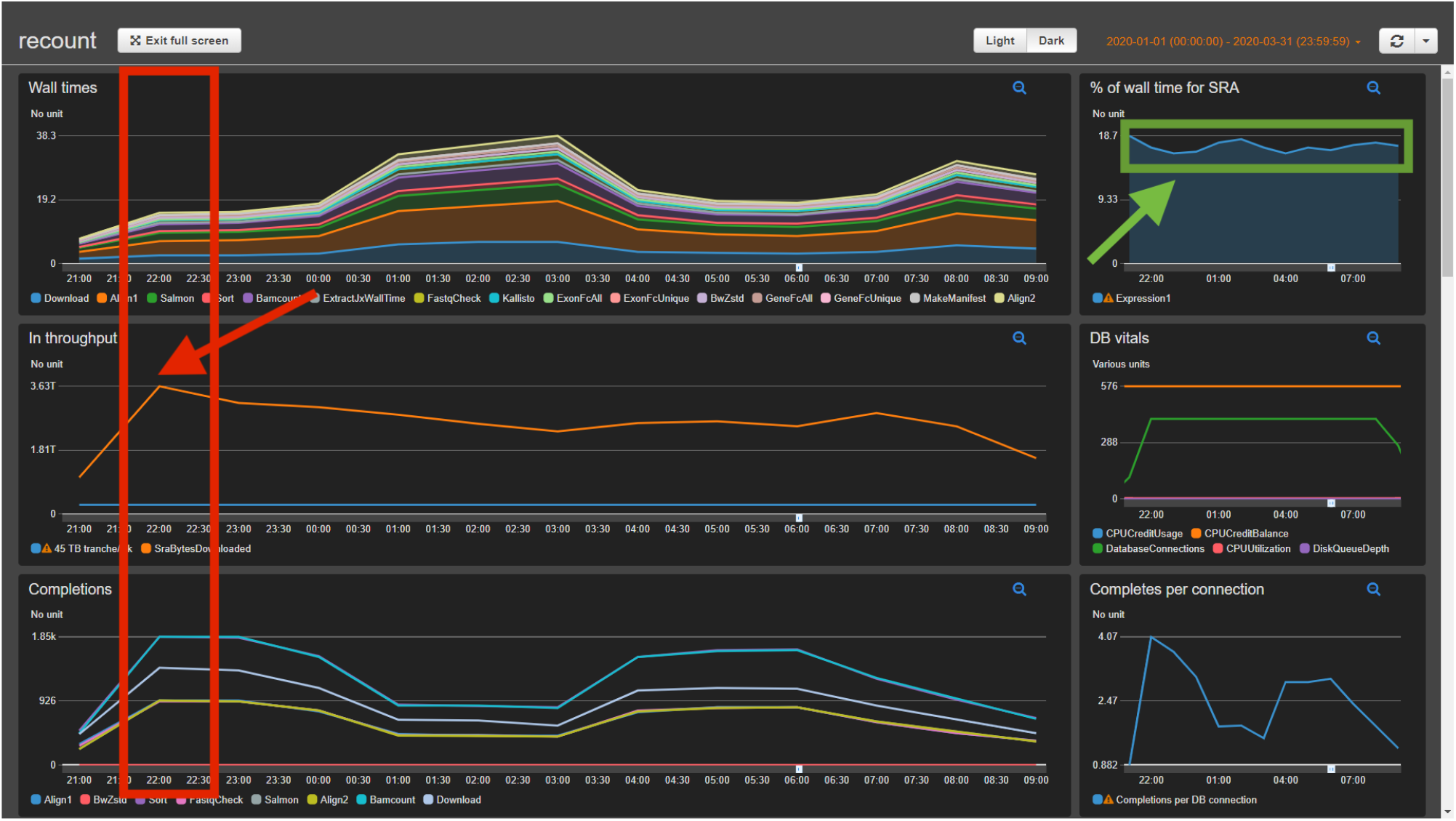
Dashboard at peak throughput. Example of how the `Monorail` dashboard captured an instance of peak throughput when using 24 out of our maximum allocated 25 simultaneous Skylake nodes on the Stampede2 supercomputer at the Texas Advanced Computer Center. The red box indicates the hour when we achieved peak throughput of around 3.63 TB per hour. The red arrow points to the data point in the “In Throughput” dashboard element showing that measurement. The green box and arrow indicate that, throughout this time period, the fraction of the wall clock time spent on the download step of the workflow stayed steady at about 15–19%, indicating that downloading from the Sequence Read Archive was not a particular bottleneck during this approximately 12-hour period. See also Supplementary Note S3.

**Supplementary Figure S6.**
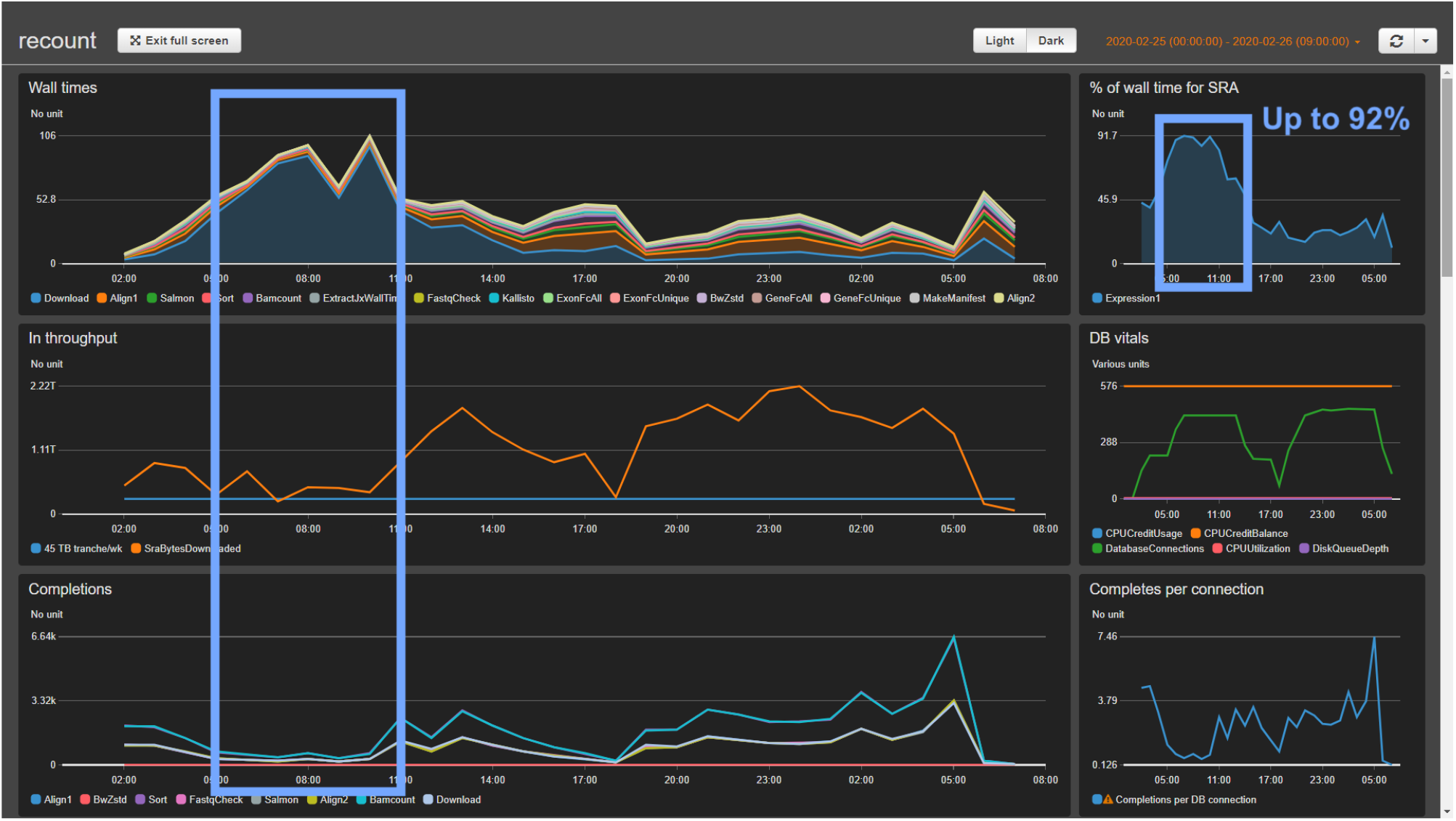
Dashboard at input bottleneck. Example of dashboard during a period where downloading from the SRA became a bottleneck. The blue rectangles highlight a period where Input Throughput was low (left panel, middle plot) and the fraction of the wall clock time spent on the download step of the workflow climbed to over 90%, staying there for a few hours before the bottleneck resolved. See also Supplementary Note S3.

**Supplementary Figure S7.**
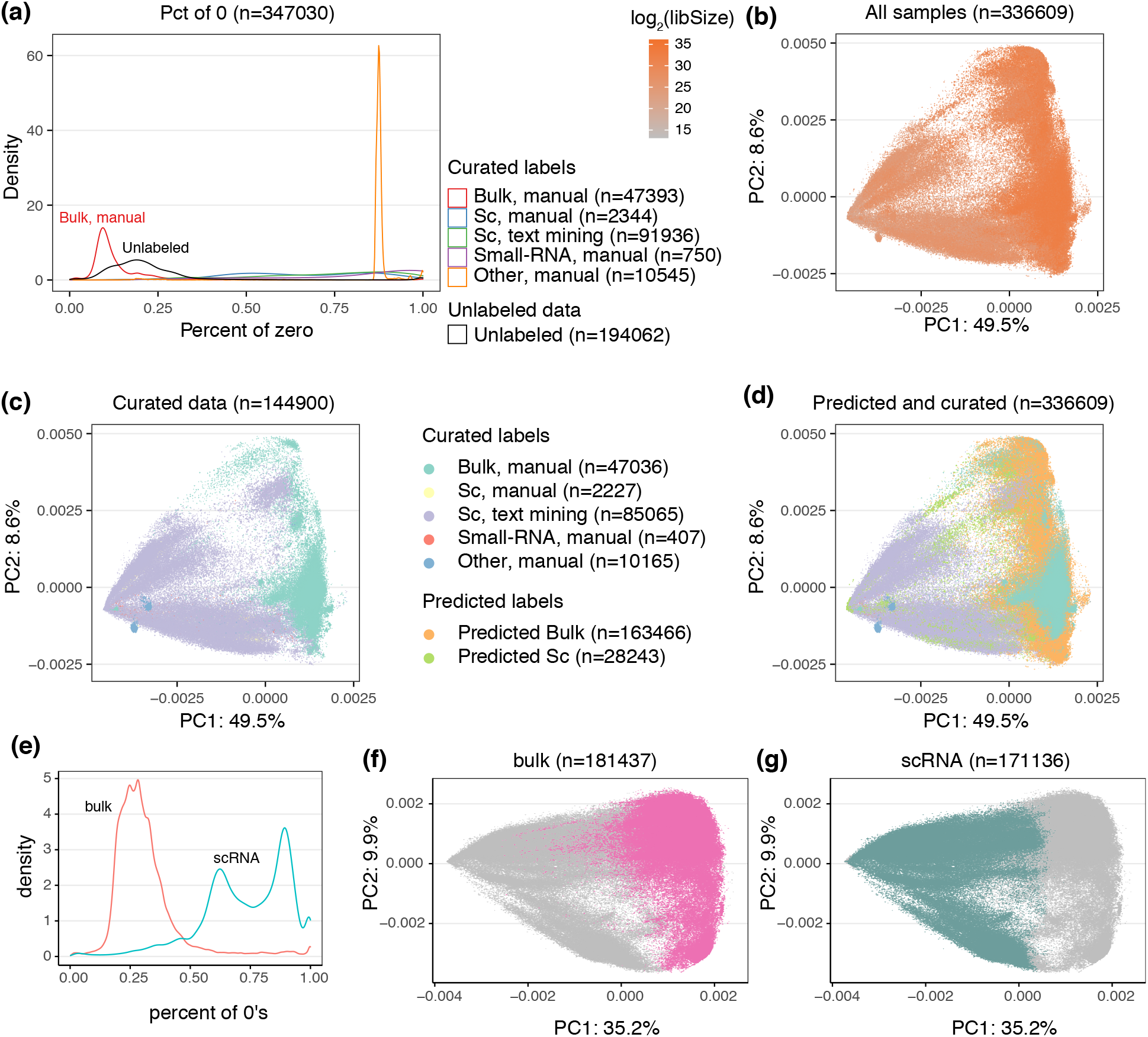
Differentiating bulk and single-cell RNA-seq data. **(a)** The distribution of zeroes for different types of labelled human data. **(b-d)** PCA of human data colored by (b) library size; (c) curated labels; (d) precited and curated labels. **(e)** Percent of zeroes from labelled mouse data. **(f-g)** Predicted labels from mouse data (f) bulk, (g) single-cell. The percentage of variance explained by the PC is given in the axis labels. Legends for (a-d): Bulk, manual - manually curated bulk samples; Sc, manual - manually curated single-cell samples; Sc, text mining - single-cell samples labelled by text analysis; Small-RNA, manual - manually curated samples which were identified as small-RNA-seq; Other, manual- manually curated samples which were identified as other assayssuch as ribosome profiling; Unlabelled - all other samples before we differentiated bulk and single-cell samples using the percentage of zero as the predictor; Predicted Bulk - predicted bulk samples using the predictor; Predicted Sc - predicted single-cell samples using the predictor. Before doing PCA, we removed some samples which we believe are not RNA sequencing samples, which accounts for the sample number difference between (a) and (b,c,d).

**Supplementary Figure S8.**
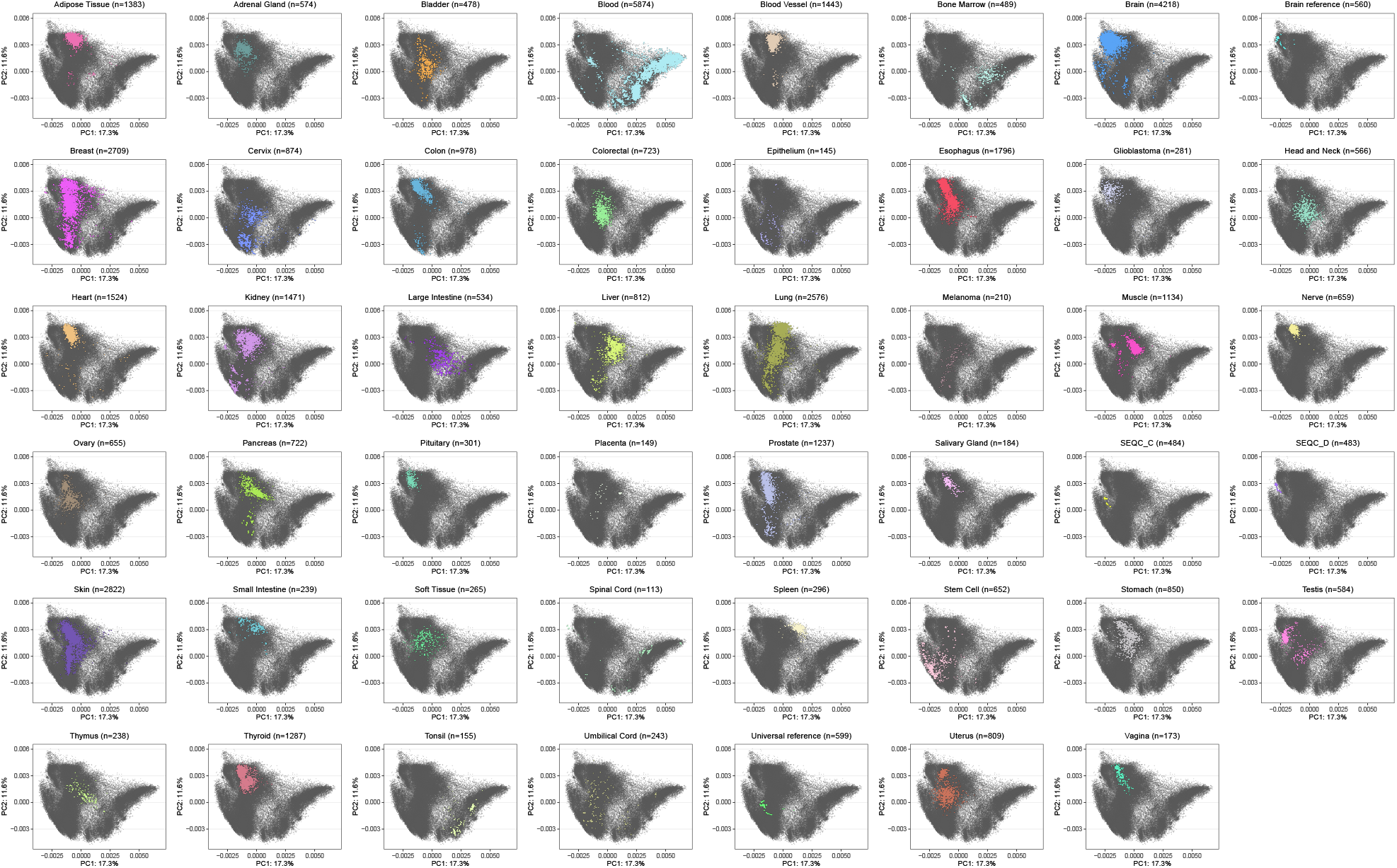
Principal component analysis of protein coding genes of bulk samples faceted by tissue types. This figure is tissue type faceted version of Figure 5. Each panel highlights one tissue type and display all other cells as gray points in the back. The percentage of variance explained by the PC is given in the axis labels.

**Supplementary Figure S9.**
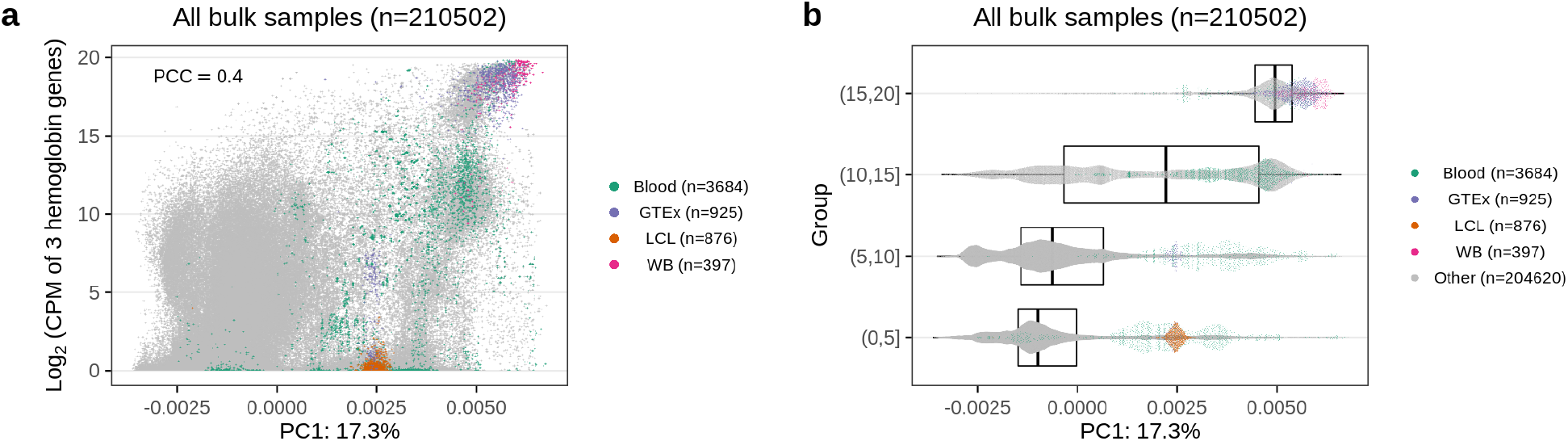
Variation of hemoglobin genes. **(a)** The sum of 3 hemoglobin genes (HBA1, HBA2, HBB) is correlated with PC1. PCC: Pearson Correlation Coefficient. Blood: manually curated blood samples; GTEx: GTEx blood samples; LCL: lymphoblastoid cell lines; WB: manually curated whole blood samples; Other: all other samples, which are colored as gray. **(b)** Similar to (a), but now we cut the *log*_2_(CPM sum) into four groups. The box shows the median, upper quartile and lower quartile of PC1 values for each group.

**Supplementary Figure S10.**
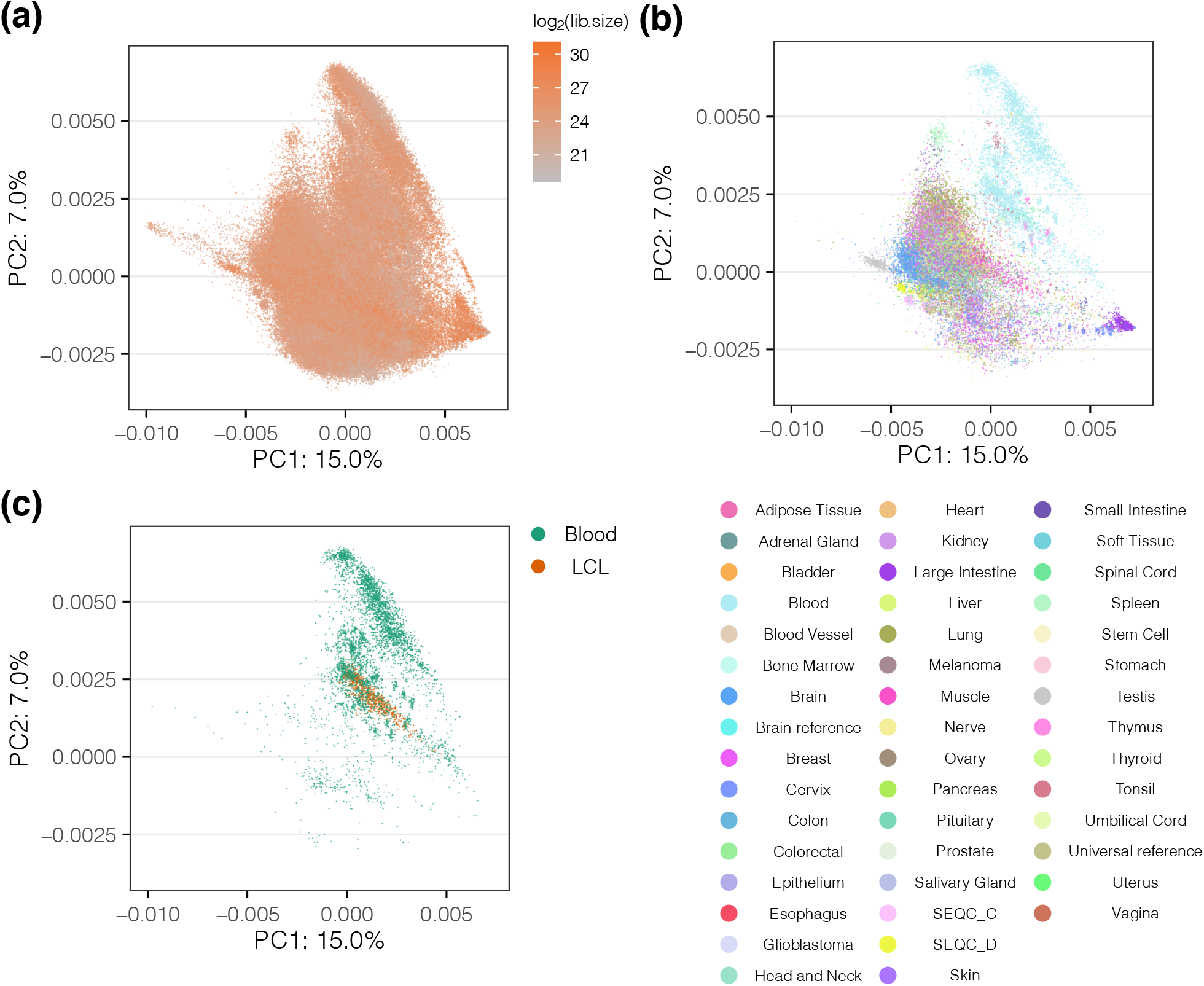
Principal component analysis of human bulk lncRNA data. This figure is similar to Figure 5, but for lncRNA data. **(a)** All human, bulk samples (207,417 samples) with color indicating the library size. **(b)** As (a) but only including samples with a labelled tissue (45,077 samples) colored by tissue. **(c)** As (a) but only including blood samples (5,966 samples) with color differentiating blood and lymphoblastoid cell lines (LCL) samples. The percentage of variance explained by the Principal Component (PC) is given in the axis labels. Note that the sample size is different to the protein coding gene data because we did separate filtering.

**Supplementary Figure S11.**
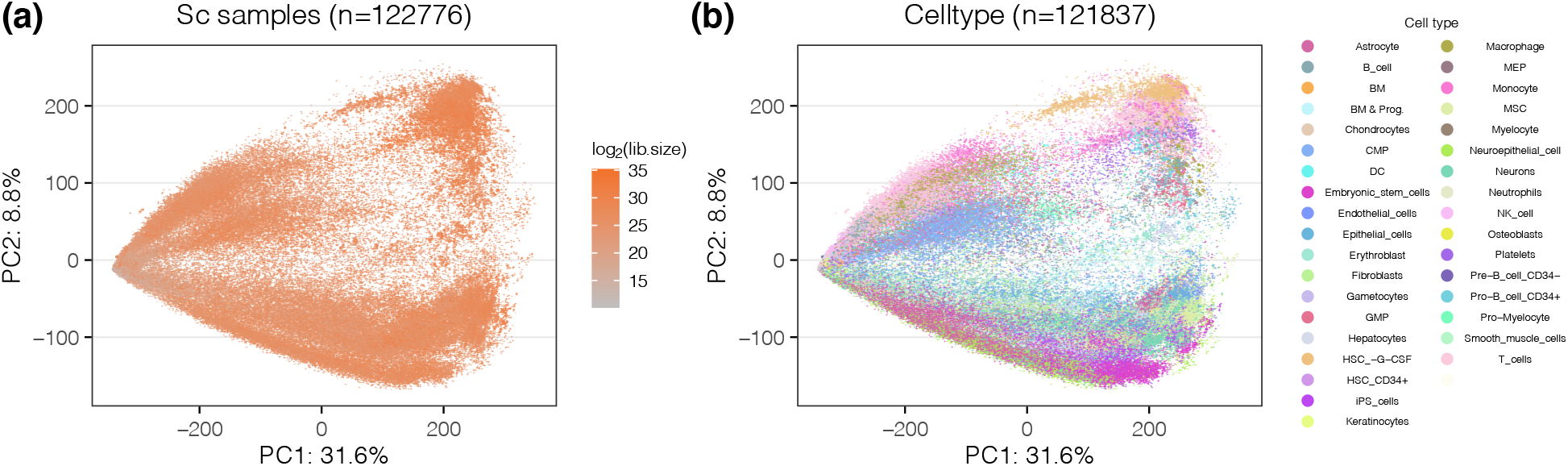
Principal component analysis of protein coding genes of single-cell samples. Similar to Figure 5, but we are now showing the top 2 PCs of protein coding genes of all human single cell samples (122,776 samples). **(a)** All human single cell samples are colored by the library size. **(b)** As (a), but only including samples with a cell type label inferred by SingleR package and colored by the cell type (121,837 samples). The percentage of variance explained by the PC is given in the axis labels.

### Supplementary Note S1

#### Detailed related work

For recount2, we previously analyzed 70,603 sequencing runs from the Sequence Read Archive (SRA), GTEx project (GTEx Consortium, 2013), and TCGA consortium (Cancer Genome Atlas Research Network et al., 2013), compiling splice-junction, gene, exon, and per-base coverages into the recount2 (Collado-Torres, Nellore, Kammers, et al., 2017) and Snaptron (Wilks et al., 2018) resources.

Other projects have worked to summarize public RNA-seq datasets, with most providing only gene- and transcript-level summaries. ARCHS4 (Lachmann, Torre, et al., 2018) used the Elysium web service (Lachmann, Z Xie, et al., 2018) – which in turn used Kallisto (Bray et al., 2016) – to quantify isoforms in 187,946 human and mouse run accessions from GEO and SRA. ARCHS4 was later updated to include over 520K accessions. The DEE2 project used STAR (Dobin and Gingeras, 2016) and Kallisto to produce gene- and transcript-level summaries for 580K run accessions, later growing to over 1 million, spanning human, mouse and seven other model organisms. Similarly, the refine.bio resource (Greene et al., 2021) used Salmon to analyze over 1.3 million samples spanning over 200 species. Tatlow et al. (Tatlow and Piccolo, 2016) used Kallisto to analyze approximately 12K TCGA and CCLE (Barretina et al., 2012) samples, also producing gene- and transcript-level summaries. Toil (Vivian et al., 2017) used STAR, Kallisto and RSEM (B Li and Dewey, 2011) to generate both spliced alignments (BAM files) and information about splice junctions detected (bedGraph files). However, it was only run on approximately 20K samples, including TCGA, TARGET, and a previous version of GTEx (about 7K samples).

Other projects have, like recount, produced larger and more multi-purpose summaries from archived RNA-seq datasets. RNASeq-er (Petryszak et al., 2017) uses the iRAP pipeline to continually analyze new RNA-seq datasets deposited in the European Nucleotide Archive. The effort has produced CRAM, bigWig and bedGraph summaries for over 1 million run accessions to date, which are accessible via a REST API. The Expression Atlas (Papatheodorou et al., 2020) draws on datasets from GEO (Barrett et al., 2013) and Array Express (Athar et al., 2019) to form a compilation of over 1M RNA assays – mostly microarray-based but also many RNA-seq – from multiple species. RNA-seq accessions are analyzed with iRAP. The Single Cell Expression Atlas (Papatheodorou et al., 2020) extends the facility to include over 100 single-cell RNA-seq studies from several species, using Alevin (Srivastava et al., 2019) for analysis.

Some design improvements in recount3 can be more directly compared and contrasted with prior projects. In particular:

- `Monorail` grid-computing design, detailed in Supplementary Note S2, is similar to that of ARCHS4 (Lachmann, Torre, et al., 2018), which also orchestrates the analysis work centrally (from a “scheduler database”) and uses containers to encapsulate the analysis pipeline. A practical difference is that `Monorail` was run on a variety of compute clusters, academic and commercial, whereas ARCHS4 was run entirely on the Amazon Web Services cloud. However, the design of ARCHS4 does allow for multi-locus computation in principle. Toil (Vivian et al., 2017), DEE2 (Ziemann et al., 2019) and the system of Tatlow and Piccolo, 2016 also use containers to encapsulate analysis.
- `Monorail`’s strategy of quantifying genes and exons starting from the bigWig coverage files, rather than from the spliced alignments, makes it particularly convenient to re-analyze datasets already summarized in `recount3` but using a new gene annotation. This is further enabled by the efficient `Megadepth` software tool (Wilks et al., 2021). This contrasts with previous projects, most of which do not make bigWig files available. Though the RNASeq-er API (Petryszak et al., 2017) does produce bigWig files, its gene- and exon-level quantifications are calculated with respect to alignments, making these far less convenient to requantify using new annotations.
- `Monorail` uses a strategy of performing full spliced alignment with respect to a reference genome, without using a gene annotation to limit or guide the alignment process. This allows recount3 to include and quantify splice junctions without any bias against unannotated junctions. This contrasts with previous projects; e.g. previous projects that used Kallisto (Bray et al., 2016) or Salmon (Patro et al., 2017) to quantify a gene annotation are not able to discover or quantify junctions absent from the annotation (Ziemann et al., 2019; Tatlow and Piccolo, 2016; Vivian et al., 2017; Greene et al., 2021). While many of the tools mentioned above do use STAR, they use it only in a mode that is guided by gene annotation (Ziemann et al., 2019; Vivian et al., 2017), and so either omit or are biased against unannotated junctions.
- recount3’s inclusion of the Snaptron infrastructure (Wilks et al., 2018) allows recount3 summaries to be rapidly queried by users. The query interface is available by REST API and via the `snapcount` R/Bioconductor package. This contrasts with past projects which have tended to make summaries available only as files — as we also do in recount3 — compiled at a pre-determined levels of granularity (study, run, etc). The Snaptron facility, by contrast, allows users to rapidly ask and answer questions about expression levels and splicing patterns across *all* run accessions in a compilation, e.g. all 316,443 human SRA samples, and retrieve a specific answer within seconds without having to download larger summaries. Importantly, these searches can be conducted not just by filtering by sample metadata, but also by specifying genomic intervals and/or expression/splicing patterns of interest. For example, a query can ask to list all of the run accessions where one splicing pattern is much more frequent than an alternative splicing pattern. Or a query can ask for all the junctions that occur more than 10 times in at least 100 run accessions. While targeted summaries are available for querying through the RNASeq-er API (Petryszak et al., 2017), these queries are less flexible, requiring the user to know a gene, study or run of interest ahead of time. A similar point is true of many of the user interfaces provided in the above systems; they usually require that the user first identify a gene, study or run of interest, before producing a figure that summarizes the corresponding data.

For details on how recount3 differs from recount2, and the advantages associated with decisions made in recount3, see Supplementary Note S6.

#### Supplementary Note S2

#### Monorail details

Here we detail the design and implementation of `Monorail`, focusing on portions of the system that generate the data summaries for recount3 and Snaptron. The system has additional tools and features not described here, but these are experimental and/or not required to produce the standard RNA-seq summaries needed for recount3.

##### Orchestration

`Monorail` follows a grid computing design, meaning that computational tasks can take place on various systems at various times, with all computation coordinated over the Internet by a few centralized services. `Monorail`’s centralized components run on Amazon Web Services. A **database server** hosts a database containing the overall data model, discussed later. This is a `db.t2.medium` instance from the Amazon Relational Database Service (RDS) running PostgreSQL version 10. A **job queue** provides a centralized, synchronized way for various analysis nodes to obtain the next available unit of work. Since it is synchronized, there is no chance of a “race condition” in the event that many analysis nodes ask for the next unit of work at the same time. This facility uses AWS’s Simple Queue Service (SQS), which also provides a degree of fault tolerance via timeout and job-visibility mechanisms. A **reference file repository** stores the reference files — e.g. genome assembly FASTA files, index files, gene annotation files — used across the project. This uses AWS’s Simple Storage Service (S3). Finally, a **centralized logging service** provides a single place for all the components of the system to keep logs. Analysis nodes, orchestration services, and client software all archive messages in this central repository. We use the AWS CloudWatch service for this facility. CloudWatch additionally allows us to visually follow the state of the system by viewing a CloudWatch Dashboard. Examples are shown in Supplementary Figures S5 & S6.

Though the orchestration layer consists of multiple cloud-based components, we adopted a decoupled web-services design that enabled local testing. This in turn allowed us to implement and iterate the design more quickly. Integration tests were accomplished using Docker Compose in conjunction with a collection of published container images that emulated each of the key components. For the PostgreSQL database server, we used the 10.4 version of the PostgreSQL published Docker image at https://hub.docker.com/_/postgres. For the SQS queueing service, we used the 0.14.6 version of the elasticmq published Docker image at https://hub.docker.com/r/softwaremill/elasticmq. For the S3 storage service, we used an adapted version of the minio published Docker image (tag 2019-03-27T22-35-21Z) at https://hub.docker.com/r/minio/minio. For testing, the CloudWatch logging service was replaced with the standard Python `logging` module.

##### Data Model

The `Monorail` data model, pictured in Figure S1, defines the kinds of information that can be tracked by the orchestration layer. For instance, the `input` **table** describes all the sequencing-read input files for all the computations. For some files, these might “point to” the dataset via an SRA accession; for others, this might use a URL to locate the file. The `annotation` **and** `source` **tables** contain information about the origin of all the reference files used, including genome indexes and gene annotations. The `analysis` table describes all the Docker and/or Singularity images that might be used to analyze an input dataset. The data model can be created and modified using Python scripts in the orchestration software, available at https://github.com/langmead-lab/recount-pump. This software uses the SQLAlchemy object-relational model to map tables in the PostgreSQL database to objects in the Python infrastructure.

##### Managers and runners

Now we describe the software that runs on the compute clusters that obtain jobs from the orchestration layer, perform the analysis, and store the outputs. The highest level, an analysis node runs a **node manager**, which launches a number of individual “job runners,” each allocated a fraction of available memory and hardware threads. The bottom layer is the **runner** which runs on the compute-node “slice” allocated to it by the node manager. A runner enters a “job loop,” where it repeatedly checks a queue of all pending tasks for the project. Once it has obtained a job, a runner launches a Singularity container that in turn runs the corresponding workflow. This is illustrated in Figure S2.

##### Workflow

The `Monorail` analysis workflow is driven by a Snakemake workflow that runs inside a Singularity container. The use of a container system allows us to package all of the constituent software tools and all their dependencies in a single image. We use Singularity (rather than Docker, for example) because it runs with non-root privileges on multi-user systems. Singularity is commonly available on scientific clusters including on our local MARCC cluster and on the Stampede2 cluster.

Here we list each major tool used in the workflow along with its version number. For a more complete account of the workflow, see the code in the `workflow/rs5` subdirectory of the https://github.com/langmead-lab/recount-pumprepository.

- Singularity: *>*=2.6.0, 3.4.2
- Snakemake: 5.4.0
- prefetch and fastq-dump (SRA Toolkit): 2.9.1-1
- parallel-fastq-dump: 0.6.3
- Google Cloud Platform gsutil: 4.47
- Rsync: 3.1.2
- Samtools: 1.9
- NCI Genomic Data Commons gdc-client: 1.3.0
- STAR: 2.7.3a
- featureCounts: 1.6.3
- Megadepth: 0.4.0
- seqtk: 1.3-r106
- GNU parallel: 20130922
- Python: 2.7.5, 3.6.7
- PyPy: 5.10.0 (Python 2.7.13)
- zstd: 1.4.0
- pigz: 2.4
- SQLite: 3.28.0
- Tabix: 1.9

##### Workflow: Alignment

The STAR aligner takes either unpaired or paired-end FASTQ files as input, together with a suffix-array-based index of the reference genome sequence. STAR then performs spliced alignment with respect to the reference sequence. Though STAR can be provided with a gene annotation, which provides “hints” as to where splice junctions might occur, Monorail does not use this feature of STAR. To emphasize: STAR can align reads in a spliced fashion across splice junctions of which it has no foreknowledge. STAR does have foreknowledge of common splice-junction motifs. Specifically, STAR understands the typical GT/AG donor/acceptor motif, as well as the and GC/AG, AT/AC motifs (and their reverse complements). Further, it allows other motifs besides these, i.e. “non-canonical” motifs. Motifs besides GT/AG receive additional penalties, with GC/AG motifs receiving a milder penalty, and AT/AC and non-canonical motifs receiving a higher penalty. More details are provided in the STAR manual.

STAR’s outputs include (a) a BAM file containing the spliced alignments, (b) a “sjout” file containing a summary of all the splice junctions detected and the level of evidence for each, (c) a file consisting of reads that failed to align, and (d) information about any chimeric alignments found. Later tools in the Monorail workflow use outputs (a)–(c) to obtain the final summary files.

##### Workflow: Gene and exon level summaries

The process by which the analysis produces gene and exon level summaries is described in Methods and illustrated in Supplementary Figure S3 and Supplementary Figure S4.

##### Workflow: Junction-level summaries

The process by which the analysis produces junctions level summaries is described in Methods and illustrated in Supplementary Figure S3 and Supplementary Figure S4.

##### Aggregation

The runners produce output files that are either transferred immediately to the aggregation node via Globus, or stored locally and periodically transferred to the aggregation node in bulk. Once a complete set of outputs are available on the aggregation node, we use our software tool (“recount-unifier”) to combine them into the tables and indexes required for recount3 and Snaptron. The unifier performs the following steps (also illustrated in Figure S4):

- Decompress exon-& junction-level summaries
- Paste exon sums together into study- and collection-level summaries
- Given gene annotations and exon sums for disjoint exonic intervals, combine these into sums across annotated genes, split by study
- Merge junction counts into a sparse matrix, only including samples which had *>* 0 splits reads for a junction
- Add information about which junctions were present in which existing gene annotations
- Split junction coverage into per-study sparse matrices in the Matrix Market format for `recount3`

The unifier output provides the underlying data files for both the Snaptron and recount3 projects, though these are formatted differently. Further, QC statistics are aggregated across sequence runs for the tranche. The unifier consists chiefly of a Snakemake workflow, using the GNU parallel utility (Tange et al., 2011) to split work across processors.

### Supplementary Note S3

#### Monorail cost & throughput

`Monorail` uses a grid computing model to spread computing work over compute clusters. Two of the clusters we used — “Stampede2” at the Texas Advanced Compute Center (TACC), and the Maryland Advanced Research Computing Center (MARCC) cluster — were free of charge, subject only to disk-space and job-queue quotas. The third venue — the Elastic Compute Cloud (EC2) on the Amazon Web Services (AWS) commercial cloud — did incur charges, allowing us to calculate the costs of at the analyses we performed there and, by extrapolation, a hypothetical overall cost for our analysis work. Here we measure and report those costs and report a few other summary measures of `Monorail`’s throughput and reliability.

Overall, we used these three clusters to analyze approximately 760,000 human and mouse sequencing runs comprising 990 TB of compressed data over six months starting October 2019 (Table 2). We used about 29,000 node hours in total, or 0.038 node-hours per sequencing run. We estimate this would cost about $0.037 per accession using equivalent cloud resources, improving on the $0.91 per accession achieved by our previous Rail-RNA system (Nellore et al., 2017).

The $0.037 per accession cost was estimated by processing a tranche of approximately 28,000 sequence runs from the SRA through our workflow, but running on EC2 instances. We chose the c5d.24xlarge instance type as it was closest in its specifications to the Skylake nodes available from Stampede2. This instance cost approximately $0.9674 per spot-instance hour in the Ohio AWS region. Note, however, that spot prices are like market prices that fluctuate according to supply and demand.

By (a) taking this spot-instance hour cost to be a cost estimate for a Stampede2 node, (b) using our measurement of the number of Stampede2 node hours (SUs or “service units”) used by our project, and (c) using the total number of run accessions we processed on Stampede2, we could extrapolate that the cost per sequencing run was approximately $0.037 per run.

While the AWS infrastructure we used for the orchestration layer and log aggregation did also incur some ongoing charges — mostly related to log aggregation (AWS CloudWatch) and queries to our data model (AWS Relational Database Service or RDS) — these charges were generally in the low hundreds of dollars per month.

When performing data analysis using the Stampede2 cluster, `Monorail` was at times able to achieve an aggregate throughput of over 3.5 terabytes (TBs) of compressed input data processed in an hour. This occurred at times when we had close to our allocated maximum of 25 simultaneous Skylake nodes scheduled. Supplementary Figure S5 shows the dashboard view at such a throughput peak; a peak throughput of approximately 3.63 TBs of input data per hour is visible in the red highlighted rectangle. This was at a time when we were using 24 out of our maximum of 25 nodes.

While we occasionally observed bottlenecks due to our download throughput from the Sequence Read Archive, evidenced by a high proportion of analysis time spent in the download step (Supplementary Figure S6), we more typically observed only about 15–19% of the time spent in the download step (Supplementary Figure S5), indicating the bottleneck was typically due to computation rather than data transfer.

When processing data on the MARCC cluster, `Monorail` achieved an aggregate throughput of approximately 0.563 compressed TBs using 11 Ivy Bridge-node compute jobs when processing runs in the TCGA project downloaded from the NCI Genomic Data Commons.

### Supplementary Note S4

#### dbGaP Access with Monorail

Most of the data processed in this project is publicly available in the Sequence Read Archive. However, both the GTEx and TCGA projects are considered protected data; potential users must go through an application process and, once access is granted, must use a key to download and decrypt the raw sequencing data. To facilitate this in `Monorail`, we support two different access methods to two protected data sources: dbGaP and the NCI Genomic Data Commons (GDC). dbGaP is the primary data source for most protected data studies’ sequence and alignment data stored in the SRA. The GDC is the primary data source for the TCGA and TARGET cancer studies’ sequence and alignment data. Since the GTEx and TCGA are both protected, we ran our entire `Monorail` analysis, including the aggregation steps, on a local high performance computing cluster, the Maryland Advanced Research Computing Cluster (MARCC).

To ensure the secure download tools have the information needed to use the decryption key (or other proof of authorization), some extra configuration is required. For dbGaP, specifically with the container approach we use in `Monorail`, the extra configuration is covered in detail here: https://github.com/langmead-lab/monorail-external/blob/master/dbgap/README.md For GDC, details are provided here: https://github.com/langmead-lab/monorail-external/blob/master/gdc/README.md

### Supplementary Note S5

#### Reference files

For the human and mouse reference genomes, we used FASTA files obtained from the iGenomes resource (https://support.illumina.com/sequencing/sequencing_software/igenome.html). For human, we used the UCSC hg38 assembly, based on GRCh38. For mouse, we used the UCSC mm10 assembly, based on GRCm38. A `STAR` index was built for each, and indexes and reference files were copied to shared filesystems on the relevant computing clusters prior to the Monorail analysis.

For both the human and mouse `STAR` indexes, we included control sequences from the ERCC (External RNA Controls Consortium, 2005) project and the synthetic spliced exons from the SIRV transcriptome project (Byrne et al., 2017). ERCC sequences were obtained from https://assets.thermofisher.com/TFS-Assets/LSG/manuals/ERCC92.zip and SIRV sequences were obtained from https://www.lexogen.com/wp-content/uploads.

When compiling gene and exon-level summaries for recount3 and Snaptron, we used multiple gene annotations. We used these human gene annotations, all compatible with the hg38/GRCh38 assembly:

- Gencode V26 (G026) (also used in GTExV8)
- Gencode V29 (G029)
- RefSeq (R109)
- FANTOM-CAT V6 (F006)

We included FANTOM-CAT (v6) (Hon et al., 2017) to be more inclusive of non-coding RNA. For QC & controls, we included the ERCC synthetic genes and SIRV transcripts.

We chose a single, recent Gencode version for our mouse annotation (M23) based on GRCm38 (mm10) in addition to the ERCC and SIRV sets mentioned above.

For both organisms, we converted a GTF of the combined set of annotations above to the GFF3 format via the `makeTxDbFromGFF` function in the `GenomicFeatures` Bioconductor package (Lawrence et al., 2013). We then passed that GFF3-formatted data to the `exonicParts` function also from the `GenomicFeatures` package setting

`linked.to.single.gene.only` to `FALSE`. This produced a set of disjoint exonic intervals covering all the annotations.

When we marked exon-exon junctions according to whether they occurred in annotation, we used a more extensive set of annotations, laid out in Table S1.

### Supplementary Note S6

#### Comparison of recount2 and recount3

recount3 includes a total of 316,443 human and 416,803 mouse samples collected from the SRA, GTEx v8 release (19,214 samples from 972 individuals and 32 tissue types), The Cancer Genome Atlas (TCGA) (11,348 samples from 10,396 individuals and 33 cancer types). This is substantially more than the 70,603 human RNA-seq samples included in our previous recount2 resource.

Also unlike recount2, roughly half the runs in recount3 are from whole-transcript single-cell protocols such as Smart-seq (Goetz and Trimarchi, 2012) and Smart-seq2 (Picelli et al., 2013).

##### Improved user interface

recount3 expands users’ options for querying the data summaries. By adding the `snapcount` Bioconductor (Huber et al., 2015) package and integrating Snaptron (Wilks et al., 2018), users can now perform rapid queries across all summaries at once, e.g. across all the 316K human SRA samples. Users can do this from the command line or from the Python or R programming languages. Between the `recount3` and `snapcount` R/Bioconductor packages, it is now much easier for users to discover relevant datasets based on metadata, to download summary data at the study or run level, and to obtain results within or across studies in metadata-rich `SummarizedExperiment` objects.

recount2 was accessible on the web through https://jhubiostatistics.shinyapps.io/recount while recount3 is now accessible through http://rna.recount.bio/, with a documentation website at http://rna.recount.bio/docs/. The recount3 website includes a study level explorer made using `shiny` (Chang et al., 2021) and `DT` (Y Xie et al., 2021) which has a selector for choosing the annotation of interest. The study selector is embedded on the documentation website such that users can now navigate the recount3 website without timeout limitations from `shinyapps.io` and is also available at https://jhubiostatistics.shinyapps.io/recount3-study-explorer/. Based on the study and annotation selection by the user, the URLs for the appropriate raw files are displayed on the study explorer. The different raw files provided by recount3 are described at http://rna.recount.bio/docs/raw-files.html. These raw files are used by `recount3` to build the `RangedSummarizedExperiment` objects on demand, unlike recount2 where the files were pre-generated. This new structure allows us to update individual components without having to re-create the final objects, reduces the required disk space, and enables building interfaces to recount3 outside of R. `recount3` uses `BiocFileCache` (Shepherd and Morgan, 2020) to cache the raw files after downloading them, thus simplifying the end-user experience that users had with `recount` (for recount2) when they repeatedly access the same data. Unlike `recount`, `recount3` does not provide a function for computing coverage for genomic regions from the recount3 bigWig files since this functionality has been greatly improved in `megadepth` (Wilks et al., 2021). Furthermore, `recount3` functions have a `recount url` argument which can be set to a different mirror or a local directory. This enables using `recount3` with data produced by `Monorail` locally that is not necessarily public.

Additionally, we provide a free, notebook-based computational resource where uses can run R and Python based analyses on the same computer cluster at Johns Hopkins University where recount3 summaries are hosted. This makes use of the existing SciServer-Compute system (Taghizadeh-Popp et al., 2020) and allows users to “bring the computing to the data” and avoid extensive downloading.

##### Improved ability to process new samples

The new, Snakemake-based (Köster and Rahmann, 2018) analysis pipeline used to produce the summaries is now much easier to use than the previous Rail-RNA software (Nellore et al., 2017). Both its analysis and aggregation components can be run from public Docker images.

##### Improved cross-study analysis

Unlike the `recount` R/Bioconductor package (for recount2), `recount3` only provides access to GTEx and TCGA data by tissue or tumor, respectively. The resulting `RangedSummarizedExperiment` objects can be combined with `cbind()` if needed. This makes it easier to access a portion of GTEx or TCGA without accessing the data from the full study.

##### Improved gene and exon quantification

Note that in recount2 we used `GenomicFeatures::exonsBy(txdb, by = “gene”)` followed by `GenomicRanges::disjoin()` (Lawrence et al., 2013) with the paramater `ignore.strand` set to the default value `FALSE` as noted in the source code of `recount::`

`reproduce ranges()` and described in the `recountWorkflow` (Collado-Torres, Nellore, and Jaffe, 2017). As described in Supplementary Note S5, in recount3 we used `GenomicFeatures::exonicParts(txdb)` (Lawrence et al., 2013). This allows `Monorail` to generate disjoint exonic intervals across genes, unlike in recount2 where the disjoint exons from one gene were not guaranteed to be disjoint exons across genes, particularly for genes from opposite strands. If exon e1 from gene g1 overlaps exon e2 from gene g2, `Monorail` generates base-pair coverage sum counts for the up to 3 disjoint exonic intervals: disjoint exonic interval A (exclusive exonic interval from exon e1), disjoint exonic interval B (shared exonic interval between exons e1 and e2), and disjoint exonic interval C (exclusive exonic internal from exon e2). This allows quantifying the expression for exons e1 and e2, and ultimately genes g1 and g2, with or without considering shared elements such as disjoint exonic interval B. The `Monorail` unifier described in Supplementary Note S2 by default does count twice disjoint exonic interval B when computing the exon e1 and e2 (once for e1 and once for e2) and ultimately the gene g1 and g2 counts, which are then accessed by `recount3`. Given file size limitations and the much larger size of recount3 compared to recount2, we did not generate additional gene and exon files excluding exonic intervals such as disjoint exonic interval B. However, using `megadepth` (Wilks et al., 2021) it is possible to re-quantify the expression of a set of samples from the recount3 bigWig files excluding disjoint exonic intervals such as disjoint exonic interval B for a given annotation of interest. Even more efficiently, `megadepth` can be used to quantify the base-pair coverage sum counts from disjoint exonic intervals such as disjoint exonic interval B and subtract it from the gene and exon counts provided by recount3. Excluding exonic intervals shared across genes, or more simply any overlapping genes, can be important for applications such as gene co-expression network inference.

As a consequence of these changes, `recount3` provides access to base-pair coverage counts aggregated at the exon level whereas `recount` (for recount2) provided access only to disjoint exons. Thus the annotation information in `recount3` for exons is much more detailed and their size is smaller and more manageable.

### Supplementary Note S7

#### recount3 R package interface

The `recount3` R package is available from Bioconductor (https://www.bioconductor.org/packages/recount3), which was first released as part of Bioconductor 3.12. This package facilitates download of recount3 data with a focus on retrieving all data from a given project or collection. Data is delivered as a standard Bioconductor data container, such as a `RangedSummarizedExperiment`, with associated metadata including metadata from the data source (ie. SRA, GTEx, TCGA) and QC data from recount3. There is additional documentation in the package vignette.

To use this package, the user has to select a project (roughly corresponding to the data from a given paper) as well as a data type. Data type is one of:

1. Gene measures (annotation dependent)
2. Exon measures (annotation dependent)
3. Junction counts (annotation independent)
4. Base-pair level data (annotation independent)

For the annotation dependent data types, an annotation needs to be selected as well. Following these choices, the data is downloaded by our data host SciServer at Johns Hopkins University (Taghizadeh-Popp et al., 2020). Data is locally cached to avoid unnecessary downloads, using the `BiocFileCache` R/Bioconductor package (Shepherd and Morgan, 2020).

Gene measures, exon measures and exon-exon junction counts are delivered as `RangedSummarizedExperiment` (RSE) objects while base-level data is delivered as collection of bigWig files.

It is not possible to retrieve parts of the data matrices, for example data on a specific gene. This is the purpose of the Snaptron web server and the `snapcount` R/Bioconductor package (Wilks et al., 2018).

`recount3` is designed to work with custom data web servers and local files, which enables researchers to use this R package with their own private data processed using the `Monorail` system.

##### Note on R package names

This manuscript describes the recount3 data collection and the `recount3` R/Bioconductor package. The recount2 data collection (Collado-Torres, Nellore, Kammers, et al., 2017) is accessed using the `recount` R/Bioconductor package, whereas the ReCount data collection (Frazee et al., 2011) does not have an associated R/Bioconductor package.

### Supplementary Note S8

#### snapcount R package interface

The `snapcount` Bioconductor package gives users an easy R interface for accessing gene, exon, and junction coverage data Snaptron queries. It replicates much of the functionality in the Snaptron Python client. This includes supports for all of the “basic” queries and filters for gene, exon and junction data. Queries can be filtered to narrow the focus to particular genes or genomic intervals, to events with certain prevalence, to events that do or don’t appear in gene annotation, or to samples with particular metadata. It also includes some higher-level queries, such as the Junction Inclusion Ratio query (JIR) for studying the relative prevalence of two or more splicing patterns, and the Tissue Specificity (TS) query for scoring junctions according to the degree to which they exhibit tissue-specific expression in the GTEx collection. Finally, it includes functions for merging results across multiple collections using the same reference genome and gene annotations (e.g. the GTEx and TCGA collections).

Data matrices returned from snapcount queries are presented to the user as dynamically generated `RangedSummarizedExperiment` (RSE) objects, which package the data together with metadata about the rows (genomic features) and columns (run accessions) of the matrix. Because the user has control over various filtering criteria pertaining to the rows and columns, this can be a more convenient and efficient way to arrive at a specific result.

The package is available at http://www.bioconductor.org/packages/snapcount.

### Supplementary Note S9

#### Selection of SRA datasets

Sequence runs from both human and mouse were selected from the Sequence Read Archive (SRA) using the search parameters defined below:

- Human Entrez search: (((((illumina[Platform]) AND rna seq[Strategy]) AND
transcriptomic[Source]) AND public[Access]) NOT
size fractionation[Selection]) AND human[Organism], run on 2019-10-06, producing 421,794 experiments to download.
- Mouse Entrez search: Uses the same search query as human, except with `mouse[Organism]`, run on 2020-01-08, producing 528,058 experiments to download.

The sequence runs for both organisms were then further filtered after retrieval of the metadata records from SRA based on the occurrence of certain single-cell RNA (scRNA) technology key words with the primary intention of removing any Droplet based technology derived sequence runs (e.g. Chromium) from the set to process through `Monorail`. The desired set of technologies was bulk-mRNA and smartSeq (and variants) due to the full transcriptome coverage profiles generated by such platforms. Droplet based technologies are typically end-biased as well as requiring substantially more effort to properly process the raw sequences due to the presence of barcodes. They are not currently supported by `Monorail`. Table S4 gives the breakdown sequence runs processed for both organisms.

We performed a text search on the retrieved data to identify single-cell samples. This was done by searching the XML files obtained from SRA for the following 20 patterns

~~~
10x cel.?seq chromium ctyoseq dronc.?seq drop.?seq fluidigm
indrop mars.?seq matq.?seq quartz.?seq sci-rna.?seq seq.?well
smart2-seq smarter smart.?seq split.?seq strt.?seq strt_seq
super.?seq
~~~

### Supplementary Note S10

#### Obtaining GTEx and TCGA data & metadata

We obtained GTEx metadata from the “Annotations” section of the GTEx portal:

https://storage.googleapis.com/gtex_analysis_v8/annotations/GTEx_Analysis_v8_Annotations_SampleAttributesDS.txt

Since GTEx includes multiple runs per aliquot, we extended the GTEx aliquot barcode with a “.#” to indicate which run the barcode is referring to. This is called “rail barcode” in our files. At the time of data collections, samples from all GTEx releases up to and including v7 were accessioned by the SRA and visible in the SRA Run Browser. Samples in GTEx v8 (excluding v6 & v7) were not accessioned in the SRA, and are not present on the SRA Run Browser. Sequence data for GTEx v7 & v8 samples (excluding v6) were available only in the AWS (v7 only) or Google Cloud Platform (GCP, v7 & v8) commercial clouds. We retrieved GTEx v7 and v8 sequence data from GCP as BAM files (9,303 files), and retrieved all the sample sequence data up to and including v6 from the SRA directly in the normal format (9,911 files). We used the “gsutil cp” tool to download from GCP, and the “prefetch” tool from the SRA-Toolkit (together with “parallel-fastq-dump”) to obtain FASTQ data from the SRA. The SRA retrieval tools are part of the `Monorail` Docker image; we do not include “gsutil” as we considered its use to be a one-time event.

Metadata for TCGA was inherited directly from recount2 (Collado-Torres, Nellore, Kammers, et al., 2017). TCGA sample sequence data was downloaded from the Genome Data Commons (GDC) using the GDC Download Client tool, version 1.4, also included in the `Monorail` Docker image.

### Supplementary Note S11

#### Quality control

We used a number of tools to collect quality-control (QC) measures for each run accession in recount3. Specifically, we used `seqtk` (H Li, 2020), the `idxstats` subcommand of `samtools`, the output of `STAR` (Dobin and Gingeras, 2016), our own `Megadepth` tool (Wilks et al., 2021), and `featureCounts` (Liao et al., 2014). The full set of QC measures are listed and briefly described at http://rna.recount.bio/docs/quality-check-fields.html.

`Monorail` runs the `seqtk fqchk` command on input FASTQ files to collect base-quality and base-composition summaries for all sequencing cycles. We distill these into a few QC measures included with every summarized run in recount3. `Monorail` uses `STAR` to align RNA-seq reads in a spliced fashion to a reference genome, without using any annotation. Files output by `STAR`, particularly the `Log.out` and `Log.final.out`, report a number of measures. We compile these into a number of QC measures, some reported separately for the two ends of a paired-end read.

The `Megadepth` tool, which runs on the BAM files output by `STAR`, also provides useful QC measures. As `Megadepth` performs bigWig conversion, it also summarizes the amount of sequencing coverage within the intervals of a provided BED file representing a gene annotation. These indicate, for example, what fraction of the coverage is within annotated genes.

Finally, `Monorail` runs `featureCounts` on the BAM files output by `STAR`. This provides a “second opinion” on the quantifications produced by `Megadepth`. We have not found compelling examples where the `Megadepth` and `featureCounts` outputs materially disagree, but we nonetheless keep both kinds of summaries as potential QC measures.

### Supplementary Note S12

#### bigWig processing with Megadepth

`Megadepth` (Wilks et al., 2021) is a new tool we built to serve two main purposes in this project:

- Efficiently convert the spliced-alignment BAM files produced by `STAR` into bigWigs files encoding the depth of coverage at each base
- Efficiently re-quantify coverage over the bigWig files produced in the previous step for a new annotation/set of intervals, avoiding the re-downloading of the original sequence + alignment steps

While there were tools available for performing each of these functions — e.g. `pyBigWig` (Ramírez et al., 2016), `WiggleTools` (Zerbino et al., 2014) — no one tool was able to do both these things efficiently in a single pass over the BAM file. `Megadepth` performs both functions efficiently using many threads, and adds no major software dependencies to `Monorail` apart from `Megadepth` itself.

Specifically, `Megadepth` produces three relevant kinds of summaries in a single pass through the `STAR` BAM file:

- Area Under Coverage (AUC)—related to mapping depth and used extensively in recount2 as described in recountWorkflow (Collado-Torres, Nellore, and Jaffe, 2017)
- Per-base coverage as a bigWig file
- Coverage across the disjoined exons from the annotations described elsewhere in this Supplement

For all coverage summaries listed above, `Megadepth` reports the number(s) for all reads mapping and those reads which mapped with minimum quality *>*= 10 separately (6 different reports).

`megadepth` also has a R/Bioconductor wrapper that allows for more easy interaction with recount3 bigWig files, as described in a separate manuscript (Wilks et al., 2021).

### Supplementary Note S13

#### recount3 data formatting

The coverage summaries provided in recount3 are stored as tab delimited matrices in GZip compressed flat files. Rows are genes or exons, and columns are samples. Coverage is stored as raw per-base counts summed over the relevant annotation interval (gene or exon). Junction files follow the Market Matrix format (*Matrix Market format* n.d.) which represents the junction coverage matrix as a sparse list of matrix coordinates for those cells which are non-0. The non-0 values represent the raw count of split reads supporting a given junction. Per-base coverage values are stored in bigWigs, one bigWig file per sample.

### Supplementary Note S14

#### Snaptron data formatting and indexing

Whereas recount3 organizes our data summaries in a form that is easy to download and use at the study level, Snaptron organizes the summaries in a way that enables queries across all data in an entire collection. Snaptron indexes the same summaries as are available in recount3, but requires a different set of indexes and data files. Snaptron stores gene, exon, and junction coverage summaries in a Snaptron-specific tab delimited sparse-matrix format. This format contains the coordinates of the gene, exon, or junction. For genes and exons, it also contains relevant information identifying the exon and/or gene. For junctions, it also contains a list of all the gene annotations it appears in. In both cases, the summary also contains a list of sample IDs where the feature occurs in, and the associated coverage (gene, exon) or number of split reads (junctions).

Snaptron also indexes a condensed form of the per-base coverage from the bigWigs for the GTEx and TCGA datasets. The SRAv3 human and SRAv1 mouse datasets are too large to fit the current per-base indexing approach in Snaptron at this time.

See the Snaptron user manual (http://snaptron.cs.jhu.edu/) and publication (Wilks et al., 2018) for details on the REST API and queries that can be made against Snaptron. Most queries can be answered within a few seconds, though this depends on the amount of data returned, server load, and network conditions between the client and server among other factors.

